# Single-cell multi-omic velocity infers dynamic and decoupled gene regulation

**DOI:** 10.1101/2021.12.13.472472

**Authors:** Chen Li, Maria Virgilio, Kathleen L. Collins, Joshua D. Welch

## Abstract

Single-cell multi-omic datasets, in which multiple molecular modalities are profiled within the same cell, provide a unique opportunity to discover the relationships between cellular epigenomic and transcriptomic changes. To realize this potential, we developed MultiVelo, a mechanistic model of gene expression that extends the RNA velocity framework to incorporate epigenomic data. MultiVelo uses a probabilistic latent variable model to estimate the switch time and rate parameters of chromatin accessibility and gene expression from single-cell data, providing a quantitative summary of the temporal relationship between epigenomic and transcriptomic changes. Incorporating chromatin accessibility data significantly improves the accuracy of cell fate prediction compared to velocity estimates from RNA only. Fitting MultiVelo on single-cell multi-omic datasets from brain, skin, and blood cells reveals two distinct classes of genes distinguished by whether chromatin closes before or after transcription ceases. Our model also identifies four types of cell states–two states in which epigenome and transcriptome are coupled and two distinct decoupled states. The parameters inferred by MultiVelo quantify the length of time for which genes occupy each of the four states, ranking genes by the degree of coupling between transcriptome and epigenome. Finally, we identify time lags between transcription factor expression and binding site accessibility and between disease-associated SNP accessibility and expression of the linked genes. We provide an open-source Python implementation of MultiVelo on PyPI and GitHub (https://github.com/welch-lab/MultiVelo).

## 1 Introduction

The regulation of gene expression from DNA to RNA to protein is a key process governing cell fates. Coordinated, stepwise gene expression changes–in which genes are turned on and off in a certain order– underlie the developmental processes by which cells specialize. Increasingly, high-throughput single-cell sequencing techniques are being applied to reveal these stepwise gene expression changes. However, because experimental measurement destroys the cell, only temporal snapshot measurements are available, and it is not possible to observe the same individual cell changing over time.

Computational approaches can leverage single-cell snapshots to infer sequential gene expression changes during developmental processes. For example, cell trajectory inference algorithms^1,2,3,4,5^ use pairwise cell similarities to map cells onto a “pseudotime” axis corresponding to predicted developmental progress. However, trajectory inference based on similarity cannot predict the directions or relative rates of cellular transitions. Methods for inferring RNA velocity^6,7^ address these limitations by fitting a system of differential equations that describes the directions and rates of transcriptional changes using spliced and unspliced transcript counts. The original RNA velocity approach^6^ relied on a steady-state assumption to fit model parameters, but later work developed a dynamical model^7^ that explicitly fits the induction and repression phases of gene expression, in addition to the steady states. Crucially, this dynamical model of RNA velocity also infers a latent time value for each cell, providing a mechanistic means of reconstructing the order of gene expression changes during cell differentiation. A recent paper further extended the RNA velocity framework to include gene expression and protein measurements from the same cells, but used the steady-state assumption to estimate parameters, and thus did not estimate latent time values for each cell^8^. Single-cell epigenome values have also been used individually to infer future directions of cell differentiation, but these approaches did not incorporate gene expression^9,10^.

Single-cell multi-omic measurements provide an opportunity to incorporate epigenomic data into mechanistic models of trancription. For example, new technologies such as SNARE-seq^11^, SHARE-seq^9^, and 10X Genomics Multiome can quantify both RNA and chromatin accessibility in the same cell. The epigenome and transcriptome both change during cellular differentiation, and thus the temporal snapshots in single-cell multi-omic datasets potentially reveal the interplay among these molecular layers. For example, if epigenomic lineage priming occurs at a particular genomic locus, single-cell multi-omic data could reveal a significant time lag between chromatin remodeling of a gene and its transcription. Similarly, observing the dynamic changes in both the expression of a transcription factor and the chromatin accessibility of putative binding sites could reveal their temporal relationship.

Existing RNA velocity models assume that the transcription rate of a gene is uniform throughout the induction phase of gene expression. However, epigenomic changes play a key role in regulating gene expression, such as tightening or loosening the chromatin compaction of promoter and enhancer regions. For example, a transition from euchromatin to heterochromatin significantly reduces the rate of transcription at that locus, because transcriptional machinery cannot access the DNA. Therefore, a more realistic model would reflect the influence of enhancer and promoter chromatin accessibility on transcription rate.

We present MultiVelo, a computational approach for inferring epigenomic regulation of gene expression from single-cell multi-omic datasets. We extend the dynamical RNA velocity model to incorporate multi-omic measurements to more accurately predict the past and future state of each cell, jointly infer the instantaneous rate of induction or repression for each modality, and determine the extent of coupling or time lag between modalities. MultiVelo uses a probabilistic latent variable model to estimate the switch time and rate parameters of gene regulation, providing a quantitative summary of the temporal relationship between epigenomic and transcriptomic changes.

We demonstrate that MultiVelo accurately recovers cell lineages and quantifies the length of priming and decoupling intervals in which chromatin accessibility and gene expression are temporarily out of sync. Our differential equation model accurately fits single-cell multi-omic datasets from embryonic mouse brain, embryonic human brain, and a newly generated dataset from human hematopoietic stem and progenitor cells. Furthermore, our model predicts two distinct mechanisms of gene expression regulation by chromatin accessibility, and we identify clear examples of both mechanisms across all of the tissues we investigated. Finally, we use MultiVelo to infer the temporal relationship between transcription factors (TFs) and their binding sites and between GWAS SNPs and their linked genes. In summary, MultiVelo provides fundamental insights into the mechanisms by which epigenomic changes regulate gene expression during cell fate transitions.

## 2 Results

### 2.1 MultiVelo: A Mechanistic Model of Gene Expression Incorporating Chromatin Accessibility

MultiVelo describes the process of gene expression as a system of three ordinary differential equations (ODEs) characterized by a set of switch time and rate parameters (Fig. 1A). The time-varying levels of chromatin accessibility (*c*), unspliced pre-mRNA (*u*), and spliced mature mRNA (*s*) are related by ODEs describing the rates of chromatin opening (*α*_*co*_) and closing (*α*_*cc*_), RNA transcription (*α*), RNA splicing (*β*), and RNA degradation or nuclear export (*γ*). We assume that chromatin opening rapidly leads to full accessibility and similarly that chromatin closing rapidly leads to full inaccessibility, a model supported by the datasets we analyzed (Fig. S3A and S3B). The single chromatin accessibility value (*c*) for a gene is calculated by summing all accessibility peaks linked to the gene; we tested multiple strategies for calculating *c* and found that they do not significantly change the results (Fig. S2). Each gene has distinct rate parameters describing its unique kinetics. We assume that the transcription rate is proportional to the chromatin accessibility *c*(*t*) and thus is time-varying, and we model the distinct phases or states *k* that a cell traverses as its time *t* advances. There are two states each for chromatin accessibility (*c*) and RNA (*u, s*): chromatin opening, chromatin closing, transcriptional induction, and transcriptional repression. Each state begins at an associated switch time (*t*_*c*_, *t*_*i*_, and *t*_*r*_; chromatin opening begins at *t*_*o*_ = 0) and converges to an associated steady state value as *t* → ∞. The rate parameters and switch times are estimated for each gene using the three-dimensional phase portrait of (*c, u, s*) triplets observed across a set of single cells. The state *k* and time *t* for each cell are determined by projecting the cell to the nearest point on the curve described by the ODEs.

**Fig. 1.**
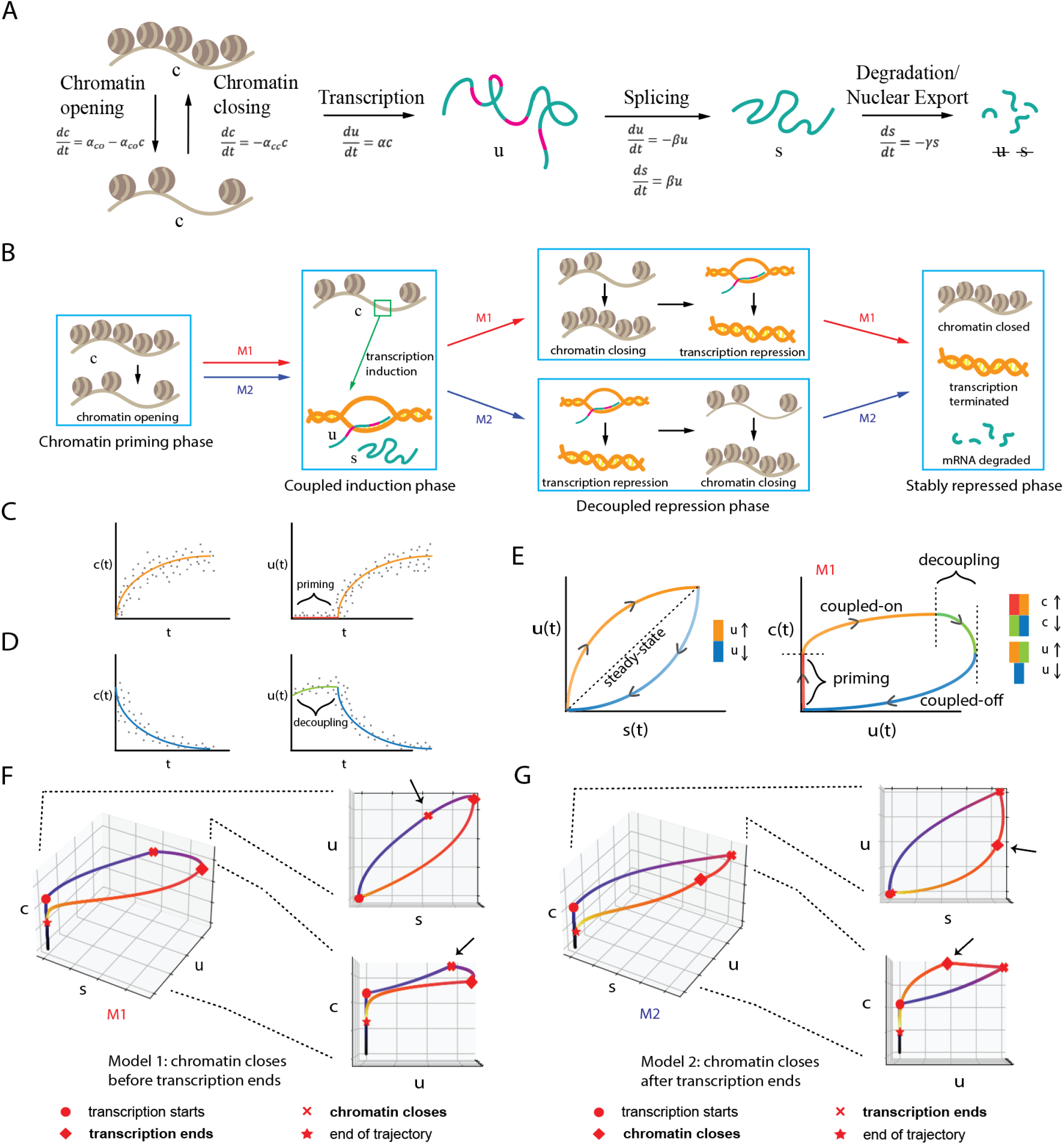
Schematic of MultiVelo approach. **A**. System of three ordinary differential equations summarizes the temporal relationship among ***c, u***, and ***s*** values during the gene expression process. **B**. Two different models (abbreviated as M1 and M2) describe two potential orderings of chromatin and RNA state changes. Chromatin accessibility starts to drop before transcriptional repression begins in M1, and the reverse happens in M2. **C**. Priming occurs when chromatin opens before transcription initiates. **D**. Decoupling occurs when chromatin closing and transcription repression begin at different times (example shown for Model 1). **E**. Phase portraits predicted by the ODE model, showing the four possible states each gene can occupy. Gene expression and chromatin accessibility are coupled in the orange and blue states, and decoupled in the red and green states. **F-G**. Simulated **(*c, u, s*)** values for a Model 1 (**F**) and a Model 2 (**G**) gene.

The mathematical formulation of the MultiVelo model immediately leads to two important insights about the relationship between chromatin accessibility and transcription during the gene expression process. First, there are multiple mathematically feasible combinations of chromatin accessibility and RNA transcription states. That is, chromatin can be either opening or closing while transcription is being either induced or repressed. This means that multiple orders of events are possible: chromatin closing can occur either before or after transcriptional repression begins (Fig. 1B). We refer to the first ordering (chromatin closing begins before transcriptional repression) as Model 1 and the second ordering as Model 2. Note that there are other mathematically possible orderings where transcription occurs before chromatin opening, but these are not biologically plausible, and we do not find convincing evidence that they occur in the datasets we analyzed (Fig. S3C).

The second insight from MultiVelo’s mathematical model is that two distinct types of discordance between chromatin accessibility and transcription can occur. At the beginning of the gene expression process, chromatin opens before transcription initiates. This creates a time interval during which *c*(*t*) is positive but *u*(*t*) and *s*(*t*) are both zero (Fig. 1C). We refer to this phenomenon as *priming*. In addition, at the end of the gene expression process, chromatin closing and transcriptional repression can occur at different times. This creates a time interval in which chromatin accessibility and gene expression move in opposite directions (Fig. 1D), a phenomenon we refer to as *decoupling*. The lengths of time during which priming and decoupling occur depend on the specific rate parameters for each gene, and thus can vary widely across genes. In between priming and decoupling intervals, when chromatin is open and transcription is active, the system converges to a steady state in which chromatin and RNA levels are coupled; similarly, when transcription is inactive and chromatin is closed, the system is in a stable repression state. These are the two stable states that differentiated cells presumably occupy most of the time.

MultiVelo infers and quantifies these phenomena of multiple orders and types of discordance through the ODE parameters estimated from single-cell data. First, the switch times (*t*_*c*_, *t*_*i*_, and *t*_*r*_) indicate when chromatin closing, transcriptional induction, and transcriptional repression begin. Thus, the lengths of priming and decoupling phases are estimated by the model: *Δt*_*priming*_ = *t*_*i*_ − *t*_*o*_ = *t*_*i*_ and *Δt*_*decoupling*_ = *t*_*r*_ − *t*_*c*_. Furthermore, because each cell is assigned latent time (*t*) and latent state (*k*) values, MultiVelo determines whether each cell is in a primed, decoupled, or coupled phase for each gene (Fig. 1E). Thus, we refer to the four possible states as *primed* (red), *coupled on* (orange), *decoupled* (green), and *coupled off* (blue). Second, the parameters fitted by MultiVelo can be used to determine, for each gene, whether its observed (*c, u, s*) values are best fit by Model 1 or Model 2 (Fig. 1F-G). Intuitively, it is possible to distinguish these models because Model 1 genes achieve their highest accessibility values during the transcriptional induction phase, while Model 2 genes reach maximum accessibility during the transcriptional repression phase (Fig. 1F-G).

### 2.2 MultiVelo Accurately Fits Simulated Data

We performed simulations to determine whether MultiVelo can recover rate parameters and switch times and distinguish Model 1 from Model 2 in the presence of noise (Fig. S1). The results indicate that MultiVelo accurately fits noisy data and can recover the underlying parameters. In addition, we found that MultiVelo distinguishes between Model 1 and Model 2 with high accuracy (98.5% of the simulated genes were correctly assigned based on model likelihood). We also confirmed that it is possible to distinguish Model 1 vs. Model 2 genes before fitting the ODE parameters by simply comparing the number of cells in the top quantiles above and below the steady-state line (95.8% of the simulated genes were correctly assigned).

### 2.3 MultiVelo Distinguishes Two Models of Gene Expression Regulation in Embryonic Mouse Brain

We first applied MultiVelo to 10X Multiome data from the embryonic mouse brain (E18). MultiVelo accurately fit the observed chromatin accessibility, unspliced pre-mRNA, and spliced mRNA counts across the population of brain cells, identifying 426 genes whose patterns fit the model with high likelihood. The resulting velocity vectors and latent time values inferred by MultiVelo accurately recover the known trajectory of mammalian cortex development. Specifically, radial glia (RG) cells in the outer subventricular zone (OSVZ) give rise to neurons, astrocytes, and oligodendrocytes^12,13,14^. Cortical layers are formed in an inside-out fashion during neuron migration with new-born cells moving to upper layers and older cells staying in deeper layers^15^. RG cells can divide into intermediate progenitor cells (IPCs) that serve as neural stem cells and further generate various mature excitatory neurons in different layers^16,17^.

Incorporating both chromatin accessibility and gene expression improves the accuracy of velocity estimation compared to RNA-only models such as scVelo (Fig. 2A). In particular, the RNA-only model predicts biologically implausible backflows inside upper layer neurons (Fig. 2B). Cell cycle scores^18,7^ indicate that the developmental process begins with a cycling population (Fig. 2C) near RG, confirming the latent time inferred by MultiVelo. MultiVelo and scVelo use similar parameter settings and estimation algorithms, suggesting that the epigenomic data provides important additional information about the past and future states of a cell, beyond what is available from transcriptomic data alone.

**Fig. 2.**
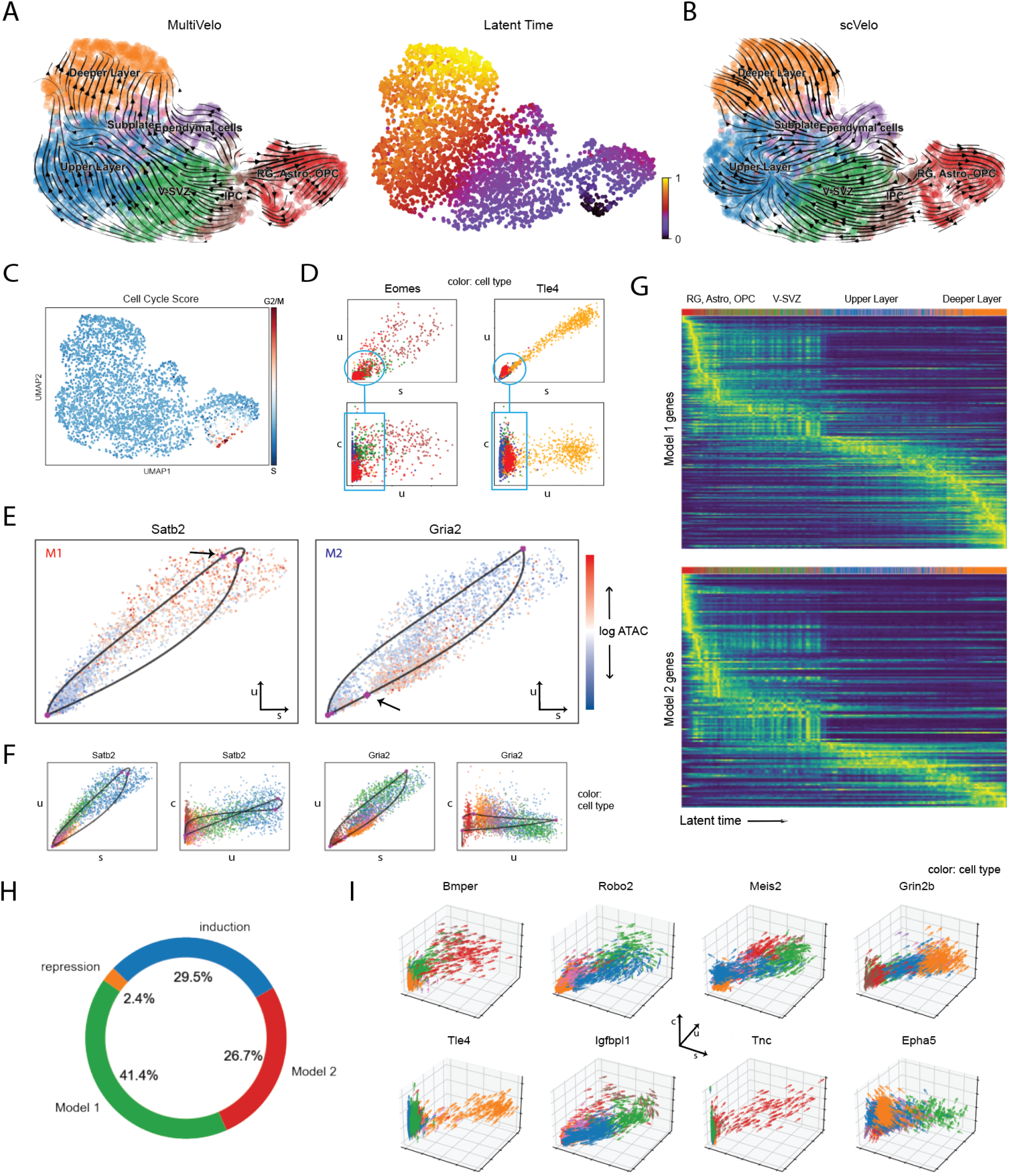
MultiVelo reveals two distinct mechanisms of gene regulation. **A**. UMAP coordinates with stream plot of velocity vectors (left) and latent time (right) from MultiVelo. **B**. Stream plot of velocity vectors estimated from RNA only by scVelo. **C**. Cell cycle score indicating active dividing and cycling population (arrow). **D**. Chromatin values better separate differentiating cells when chromatin opening precedes transcription. **E**. RNA phase portraits (*u* vs. *s*) colored by *c* values show clear differences between Model 1 (left) and Model 2 (right) genes. **F**. Additional phase portraits for the genes shown in **E. G**. Heatmaps of Model 1 and Model 2 gene expressions as a function of latent time. Color represents smoothed spliced counts. Model 2 genes tend to achieve highest expression earlier in latent time than Model 1 genes. **H**. Relative proportion of each type of kinetics across all fitted genes (n=865). Note that genes with partial kinetics (induction-only or repression-only) cannot be identified as Model 1 or Model 2. **I**. MultiVelo predicts 3D velocity vectors, which can be visualized as three-dimensional arrow plots.

We expect the addition of chromatin accessibility to be most helpful for distinguishing cell states where chromatin remodeling and gene expression are out of sync, such as when a gene’s promoters and enhancers have begun to open but little transcription has occured. Two clear examples are *Eomes* and *Tle4*, canonical markers of IPCs and deep layer neurons^19,20,21,22^. RNA transcripts from these genes are highly expressed in only one or two specific cell types. The remaining cells are densely clustered near the origin of the (*u, s*) phase portrait, making it difficult for RNA velocity methods to distinguish their relative order (Fig. 2D). However, the chromatin accessibility of these genes begins to rise before the gene expression, revealing gradual changes that are not visible from gene expression alone. To put it another way, incorporating chromatin allows us to infer 3D velocity vectors indicating each cell’s predicted differentiation for each gene, better resolving cellular differences than the 2D phase portraits from RNA alone.

MultiVelo identifies clear examples of genes that are best described by either Model 1 and Model 2 in this dataset. Comparing the phase portraits of the genes assigned to Model 1 and Model 2 shows clear differences in the timing of maximum chromatin accessibility, consistent with the model predictions (Fig. 2E). Model 1 genes such as *Satb2* reach maximum chromatin accessibility during the transcriptional induction phase (above the diagonal steady-state line on the phase portrait^6^), while the accessibility of Model 2 genes like *Gria2* is highest during the transcriptional repression phase (below the diagonal steady-state line). The distinction between Model 1 and Model 2 is also evident when inspecting pairwise phase portraits of *c, u* and *c, s* (Fig. 2F). However, the models cannot be distinguished by inspecting the RNA information alone in a phase portrait of *u, s*; the distinction requires the additional information from chromatin.

We further investigated the Model 1 and Model 2 genes to see if they have any characteristic properties. Gene ontology (GO) analysis showed that M2 genes are significantly enriched for terms related to the cell cycle, such as “positive regulation of cell cycle”, “mitotic cell cycle”, and “regulation of cell cycle phase transition”. Furthermore, Model 2 genes tend to achieve their highest spliced expression earlier in latent time than Model 1 genes (*p* = 9× 10^−7^, Wilcoxon rank-sum one-sided test; Fig. 2G). We hypothesize that cells may use Model 2 for rapid, transient activation of genes that do not need to maintain expression, whereas Model 1 may be useful for genes that need to be stably expressed.

We next looked at how often each type of gene expression kinetics (induction only, repression only, Model 1, or Model 2) occurred. Most of the highly variable genes show both induction and repression phases (a complete trajectory), and for genes that only have partial trajectories, induction-only phase portraits appear more often than repression-only (29.5% vs 2.4% of variable genes; Fig. 2H). Note that, because Model 1 and Model 2 make the same predictions during the induction phase, we cannot distinguish Model 1 vs. Model 2 for induction-only genes. Among the genes with both an induction and repression phase, the majority are best explained by Model 1 (41.4% of variable genes), while the remainder are best fit by Model 2 (26.7% of variable genes). The fact that Model 1 is more common is consistent with the expectation that chromatin state changes generally precede mRNA expression changes.

Whether genes have complete or partial kinetics, MultiVelo fits ODE parameters that describe the three dimensional trajectory of their chromatin accessibility and gene expression dynamics (Fig. 2I). By modeling a time-varying transcription rate, MultiVelo is able to better capture the different types of curvatures in the RNA phase portraits (Fig. S4B), whereas the RNA-only model cannot capture such curvature differences^23^. Genes with different model assignments and kinetics do not show significant differences in likelihood or total counts, indicating that technical artifacts do not account for the phenomena (Fig. S4C).

### 2.4 MultiVelo Identifies Epigenomic Priming and Decoupling in Embryonic Mouse Brain

An exciting property of MultiVelo is its ability to quantify the discordance and concordance between chromatin accessibility and gene expression within differentiating cells. Specifically, MultiVelo infers switch time parameters that identify the intervals during which each gene is in one of the four possible states (primed, coupled on, decoupled, and coupled off; see Fig. 1E). We next investigated whether these inferred states and time intervals can accurately capture the interplay between epigenomic and transcriptomic changes in embryonic mouse brain cells.

MultiVelo identifies clear examples of each of the four states in the 10X Multiome data (Fig. 3A). For example, *Grin2b* is an induction-only gene with expression increasing toward the neuronal fate, so only induction states–primed and coupled on–were predicted for this gene (Fig. 3A, left). The phase portrait of *Nfix*, a Model 1 gene, possesses a complete trajectory shape and was labeled with all four states (Fig. 3A, middle). Conversely, *Epha5* is a Model 2 gene, and its accessibility continues to rise throughout the whole time range without an observed closing phase, so it only occupies the coupled on and decoupled states (Fig. 3A, right).

**Fig. 3.**
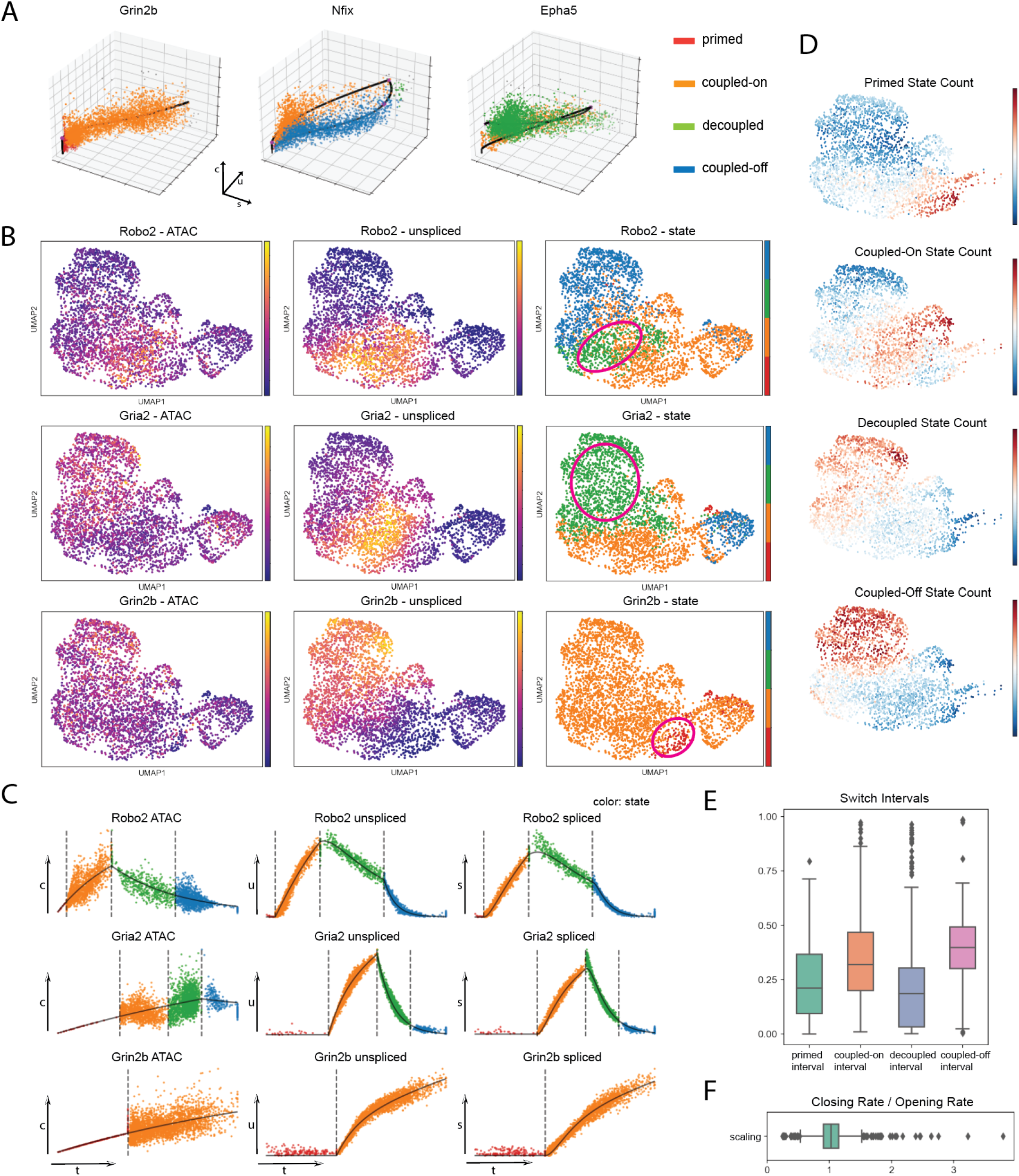
MultiVelo quantifies epigenomic priming and decoupling in embryonic mouse brain. **A**. 3D phase portraits overlaid with MultiVelo fits (solid lines) and inferred states (colors). Each point represents the (*c, u, s*) values observed for one gene in one cell. **B**. UMAP plots colored by *c* (**Left**), *u* (**Middle**, and state assignments (**Right**) for genes predicted by MultiVelo to have significant priming or decoupling intervals. Regions with priming or decoupling are circled. **C**. Observed values for *c* (**Left**), *u* (**Middle**) and *s* (**Right**) plotted as a function of latent time and colored by state assignment. Vertical lines indicate inferred switch times. **D**. UMAP plots colored by the number of genes in each cell assigned to each of the four states. **E**. Box plots summarizing the lengths of each of the four states across all fitted genes. **F**. Box plot summarizing the ratio between chromatin closing rate *α*_*cc*_ and opening rate *α*_*co*_ across all fitted genes.

The state assignments can be confirmed qualitatively by plotting accessibility (*c*) and expression (*u* and *s*) on UMAP coordinates and examining them side-by-side (Fig. 3B). Visually, we observe that the colors of the *c* and *u* UMAP plots match when the state assignments are coupled on or coupled off, and the differences in color occur when the assigned states are primed or decoupled. For example, the largest discrepancy between *Robo2* RNA expression and chromatin accessibility occurs in the circled region, which is predicted to be in the decoupled state (Fig. 3B, top). *Robo2* is a Model 1 gene; after chromatin closing begins, expression stays at a relatively high level, even though its accessibility has already experienced a drop toward the maturing neurons. Similarly, the accessibility of *Gria2* differs from RNA in the decoupled state (Fig. 3B, middle). The chromatin accessibility of *Gria2*, a Model 2 gene, continues to increase beyond the transcriptional induction phase. Furthermore, the gene *Grin2b* shows a clear example of the chromatin priming phase, during which chromatin opens prior to RNA production (Fig. 3B, bottom).

Plotting *c, u*, and *s* along the inferred time *t* for each gene allows us to inspect the state transitions in detail (Fig. 3C). First, the *u*(*t*) and *s*(*t*) values for *Robo2* show two inflection points during the transcriptional repression phase, corresponding to the transitions from coupled on to decoupled states and from decoupled to coupled off states (Fig. 3C, top). This pattern suggests that the distinct effects of chromatin closing and transcriptional repression are visible in *u*(*t*) and *s*(*t*). In other words, MultiVelo predicts that for *Robo2*, chromatin closing decreases the overall transcription rate as RNA level begins to drop immediately following the chromatin switch. The subsequent switch of transcription rate from positive to zero causes a second inflection, leading to even more rapid down-regulation of RNA expression. The plots of *c*(*t*), *u*(*t*), and *s*(*t*) for *Gria2* show the opposite trend: *c* continues to rise even after the switch to transcriptional repression, causing *c* and *u* to move in opposite directions during the decoupled state (Fig. 3C, middle). In *Grin2b*’s long priming phase, *c*(*t*) begins to rise while *u*(*t*) and *s*(*t*) stay at zero (Fig. 3C, bottom).

Because MultiVelo fits rate and switch time parameters for each gene, our analysis provides an opportunity to observe general trends in gene regulation. First, to determine whether the states of different genes are temporally coordinated, we counted the number of high-likelihood genes in each state per cell. There is indeed a cascade of state transitions through the neuronal clusters; multiple genes per cell are often simultaneously in the priming or decoupling states (Fig. 3D). Second, we looked for trends in the switch time and rate parameters. We placed each gene’s induction/repression cycle on a time scale between 0 and 1 and found that the coupled on and coupled off states account for a larger proportion of the gene expression process than the primed and decoupled states (Fig. 3E). This makes sense, because even if genes experience some level of decoupling and time lag between the two modalities, chromatin accessibility and gene expression should still be generally correlated^24,25,26,27^. The median primed interval length is 21% of the overall time, and the median decoupled interval length is 19% of the overall time. Furthermore, we can rank genes by how long their priming and decoupling intervals are to find examples of discordance between accessibility and expression (Fig. S4D). Additionally, we found that chromatin generally opens and closes at similar rates: the median ratio between inferred chromatin closing rate (*α*_*cc*_) and chromatin opening rate (*α*_*co*_) is almost exactly 1 (Fig. 3F).

### 2.5 MultiVelo Quantifies Epigenomic Priming in SHARE-seq Data from Mouse Hair Follicle

A recent study^9^ used SHARE-seq to investigate the rapid proliferation of transit-amplifying cells (TAC) in hair follicle tissue, which give rise to several mature effector cells, including inner root sheath (IRS) and layers of hair shaft: cuticle, cortical layer, and medulla^28^. When applied to this dataset, MultiVelo correctly identified direction of differentiation from TACs to IRS and hair shaft cells (Fig. 4A), consistent with the diffusion map^29^ analysis reported in the initial paper^9^. Latent time predicted the TACs to be the root cells–agreeing with biological expectation–whereas velocity analysis using RNA alone failed to capture the hair-shaft differentiation direction (Fig. 4B). We observed significantly more induction-only and fewer Model 2 genes in this dataset compared to mouse brain (Fig. 4C).

**Fig. 4.**
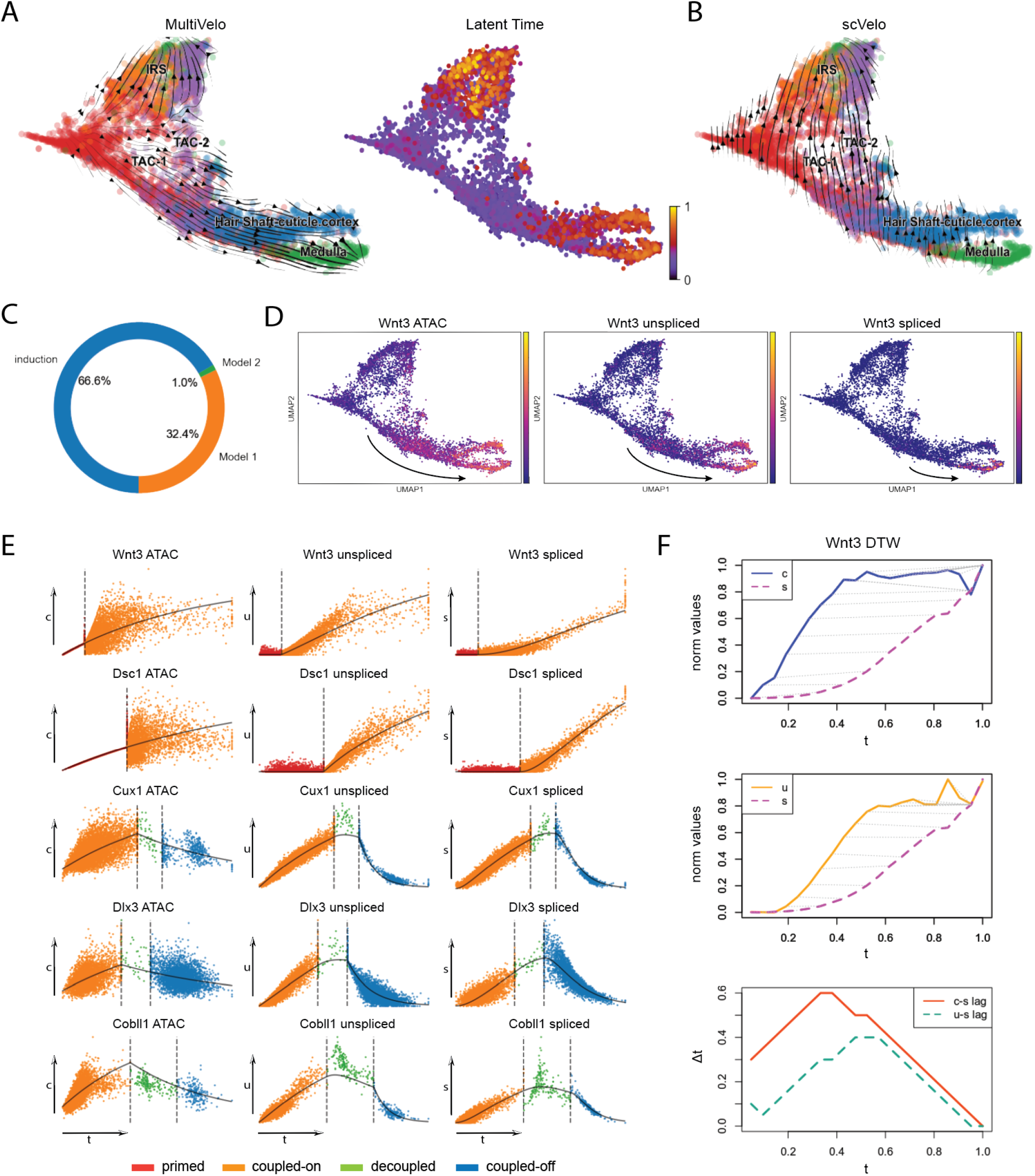
MultiVelo quantifies epigenomic priming in mouse skin. **A**. UMAP coordinates with stream plot of velocity vectors (**Left**) and latent time (**Right**:) from MultiVelo. **B**. Velocity streamplot from RNA-only model (scVelo). **C**. Relative proportion of each type of kinetics across all fitted genes (n=960). **D**. UMAP coordinates colored by *c* (**Left**), *u* (**Middle**) and *s* (**Right**) values for *Wnt3*. **E**. Examples of genes showing priming or decoupling. Observed *c* (**Left**), *u* (**Middle**) and *s* (**Right**) values plotted as a function of latent time and colored by state assignment. Vertical lines indicate inferred switch times. **F**. Dynamic time warping alignment of *c* and *s* values (**Top**) and *u* and *s* values (**Middle**) for*Wnt3*. Dotted gray lines indicate corresponding time points after alignment. **Bottom**: instantaneous time lags computed by subtracting times of aligned time points from the previous two panels.

One of the key results of the original SHARE-seq paper was the identification of genes where promoter and enhancer chromatin accessibility presaged gene expression, a phenomenon the authors termed “chromatin potential”. The clearest example of this phenomenon was *Wnt3*, which encodes a paracrine signaling molecule and is important in controlling hair growth^30^. Indeed, UMAP plots colored by accessibility, and unspliced and spliced mRNA expression show a clear time delay across modalities (Fig. 4D). We next examined the other genes identified in the SHARE-seq paper. Our fitted models show that MultiVelo faithfully captured the dynamics of each gene and provide clear illustrations of priming and decoupling regions (Fig. 4E). For instance, *Wnt3* and *Dsc1* show induction-only patterns and a priming state at the beginning while *Cux1, Dlx3*, and *Cobll1* have both induction and repression states with a short decoupling period in the middle.

To further quantify the temporal relationship between accessibility, unspliced expression, and spliced expression, we used dynamic time warping (DTW)^31^ to align the time series values for each molecular layer. DTW nonlinearly warps two time series to maximize their similarity and identify possible lagged correlation. DTW results on *Wnt3* show that the optimal warping function maps each point on the *c* time series forward in time, consistent with chromatin accessibility preceding gene expression (Fig. 4F, top). Unspliced and spliced expression show a similar pattern but with a shorter time delay (Fig. 4F, middle). Because DTW maps each time point on the earlier curve to a time point on the later curve, the time lag at each point in time can be computed by subtracting the times of the matched points (Fig. 4F, bottom). This analysis shows that both the delay between *c* and *s* and the delay between *u* and *s* remain positive throughout the observed time. In addition, the delay between *c* and *s* is longer than the delay between *u* and *s* throughout the observed range, with the maximum *c* and *s* delay reaching 0.6 (out of a total time range of 1).

### 2.6 MultiVelo Reveals Early Epigenomic and Transcriptomic Changes in Human Hematopoietic Stem and Progenitor Cells

Hematopoietic progenitors consist of stem-like cell populations that rapidly and continuously differentiate into various intermediate and mature blood cell types with progressively reduced self-renewal potential as they enter more lineage-restricted states^32,26^.

We cultured purified human CD34+ cells for 7 days, then sequenced them using the 10X Multiome platform. We obtained 11,605 high-quality cells post-filtering with both single-nucleus RNA-seq and ATAC-seq data. Using previously described marker genes^33,34,35,36^, we identified clusters resembling many of the populations of early blood development (Fig. S5A), including HSCs, multi-potent progenitors (MPP), lymphoid-primed multipotent progenitors (LMPP), granulocyte-macrophage progenitors (GMP), and megakaryocyte-erythrocyte progenitors (MEP). We also identified clusters resembling early granulocytes, erythrocytes, dendritic cells (DC), and platelets.

Blood cell differentiation is a challenging system to model with RNA velocity^23^, but we find that incorporating chromatin information significantly improves the local consistency and biological accuracy of predicted cell directions (Fig. 5A). In comparison, velocity vectors inferred from RNA alone do not accurately reflect the known differentiation hierarchy of HSPCs. As with the mouse brain, MultiVelo predicts Model 1 to be more common than Model 2 in this dataset; induction-only is the third most common gene class (Fig. 5B). The median lengths of observed primed and decoupled intervals are shorter than those of the coupled phases (Fig. 5C). These patterns are consistent with what we observed in the mouse brain dataset, suggesting a possible common underlying biological mechanism.

**Fig. 5.**
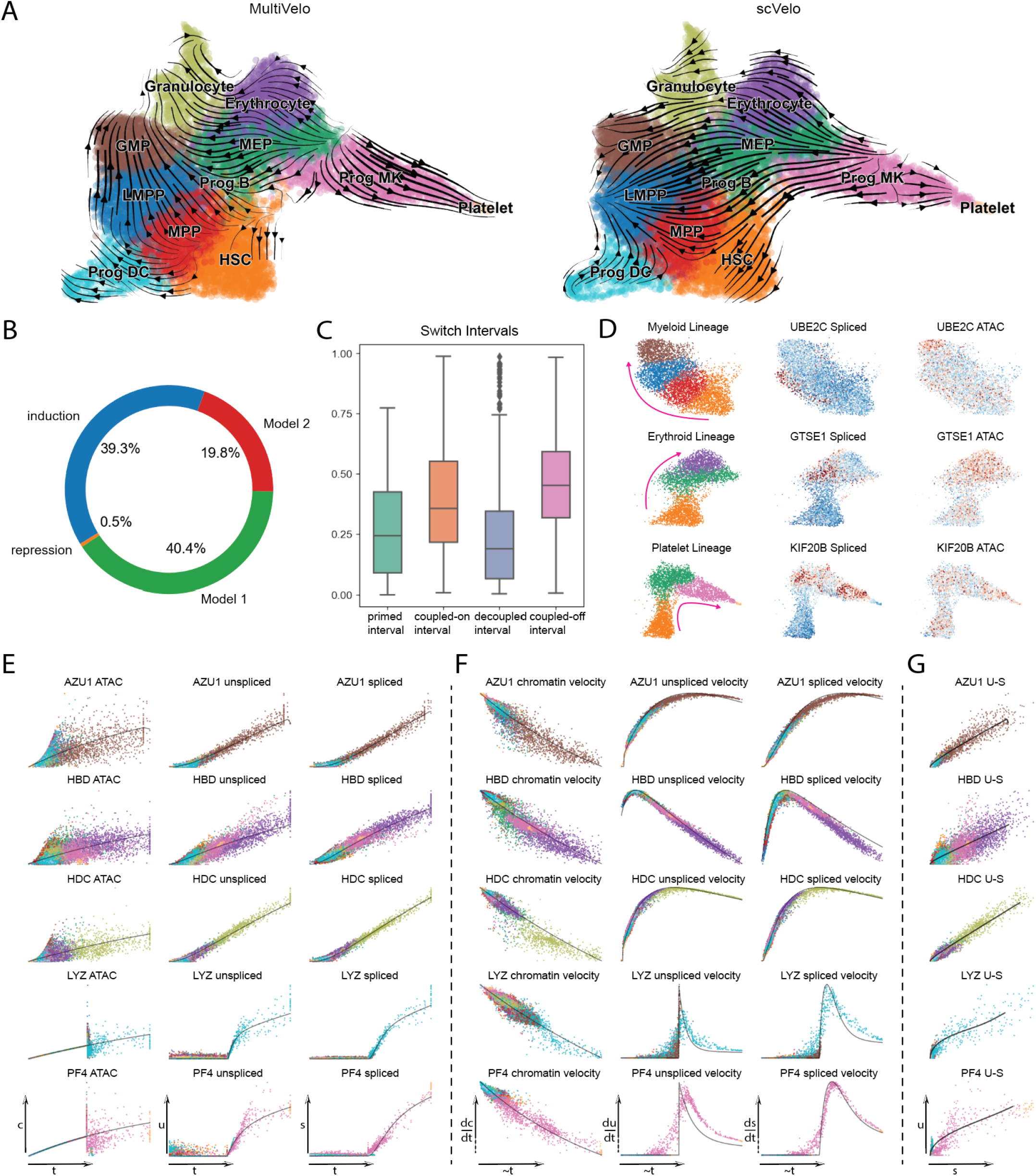
MultiVelo identifies priming in hematopoietic stem cells. **A**. UMAP coordinates with stream plot of velocity vectors inferred by MultiVelo (**Left**) and an RNA-only model (scVelo). Cell types were annotated based on marker gene expression (Fig. S5A). **B**. Relative proportion of each type of kinetics across all fitted genes (n=936). **C**. Box plots summarizing the lengths of each of the four states across all fitted genes. **D**. Several G2/M cell cycle phase markers show Model 2 expression pattern towards different lineages. **E**. Examples of genes showing priming or decoupling. Observed ***c, u***, and ***s*** values plotted as a function of latent time and colored by cell type. **F**. Corresponding velocity vectors of the same genes as in **E**. Cell velocities and times have been smoothed by RNA neighbors. Note that all velocity values are non-negative, and the lowest velocities are not necessarily at 0. **G**: RNA phase portraits of the same genes as in E-F.

As with the mouse brain dataset, Model 2 genes in the HSPC dataset are significantly enriched for GO terms related to the cell cycle. The terms “regulation of mitotic cell cycle”, “regulation of mitotic metaphase/anaphase transition”, and “regulation of mitotic sister chromatid separation” are all enriched in Model 2 genes at FDR < 0.002. If we examine the separate trajectories toward myeloid, erythroid, and platelet lineages, many G2/M phase marker genes^18^ show clear Model 2 patterns, with highest chromatin accessibility after expression begins to drop (examples shown in Fig. 5D).

We further investigated whether Model 1 and Model 2 genes differ in their histone modification profiles. Because classically defined subpopulations of HSPCs can be sorted using FACS, bulk ChIP-seq data are available for some of the cell subsets in our analysis. Using these bulk datasets^37^, we compared the levels of H3K4me3, H3K4me1, and H3K27ac in FACS-purified HSCs at chromatin accessibility peaks linked to Model 1 vs. Model 2 genes (Fig. S5C). We found that Model 2 genes show significantly higher H3K4me3 (*p* = 0.016, one-sided Wilcoxon rank-sum test), a mark of active promoters. In contrast, Model 1 genes show somewhat higher H3K4me1 (*p* = 0.097), a primed enhancer mark. Both models show similar H3K27ac (an active enhancer marker) (*p* = 0.48) in HSCs.

The gene models fit by MultiVelo reveal many examples of priming (Fig. 5E). Several terminal cell-type specific markers show induction-only dynamics with an increase in chromatin accessibility followed by increasing gene expression (*AZU1* in GMP, *HBD* in erythrocytes, *HDC* in granulocytes, *LYZ* in DC progenitors, and *PF4* in the megakaryocyte (MK) progenitors direction)^38,36^. In HSPCs, we again see some clear examples of long priming periods, such as in *LYZ* and *PF4*.

Plotting velocities allows us to examine local chromatin and RNA trends in more detail (Fig. 5F). While the chromatin shows most potential (highest velocity) at the beginning for these genes, for RNA, stem cell populations such as HSC, MPP, MEP, and GMP show increased potential during their differentiation process towards one lineage. More differentiated cell types lose the ability to maintain such potential and gradually approach equilibrium (zero velocity), even though expression is still increasing somewhat. Note that even though the overall expression elevates, and velocities stay positive, local acceleration can still switch signs. MultiVelo is able to capture such rich information about the direction and rate of differentiation due to the joint mathematical modeling of chromatin and mRNA. Adding the chromatin significantly enriches the information available from RNA, as can be seen by inspecting RNA-only phase portraits (Fig. 5G).

### 2.7 MultiVelo Relates Transcription Factors, Polymorphic Sites, and Gene Expression in Developing Human Brain

We next applied MultiVelo to a recently published 10X Multiome dataset from developing human cortex^39^. As with the embryonic mouse brain dataset, MultiVelo inferred velocity vectors consistent with known patterns of brain cell development (Fig. 6A). MultiVelo correctly inferred a cycling population of cells near radial glia as the cell type earliest in latent time. In contrast, velocity vectors inferred without chromatin information predicted incongruous backflows in intermediate progenitor cells and upper layer excitatory neurons (Fig. 6B).

**Fig. 6.**
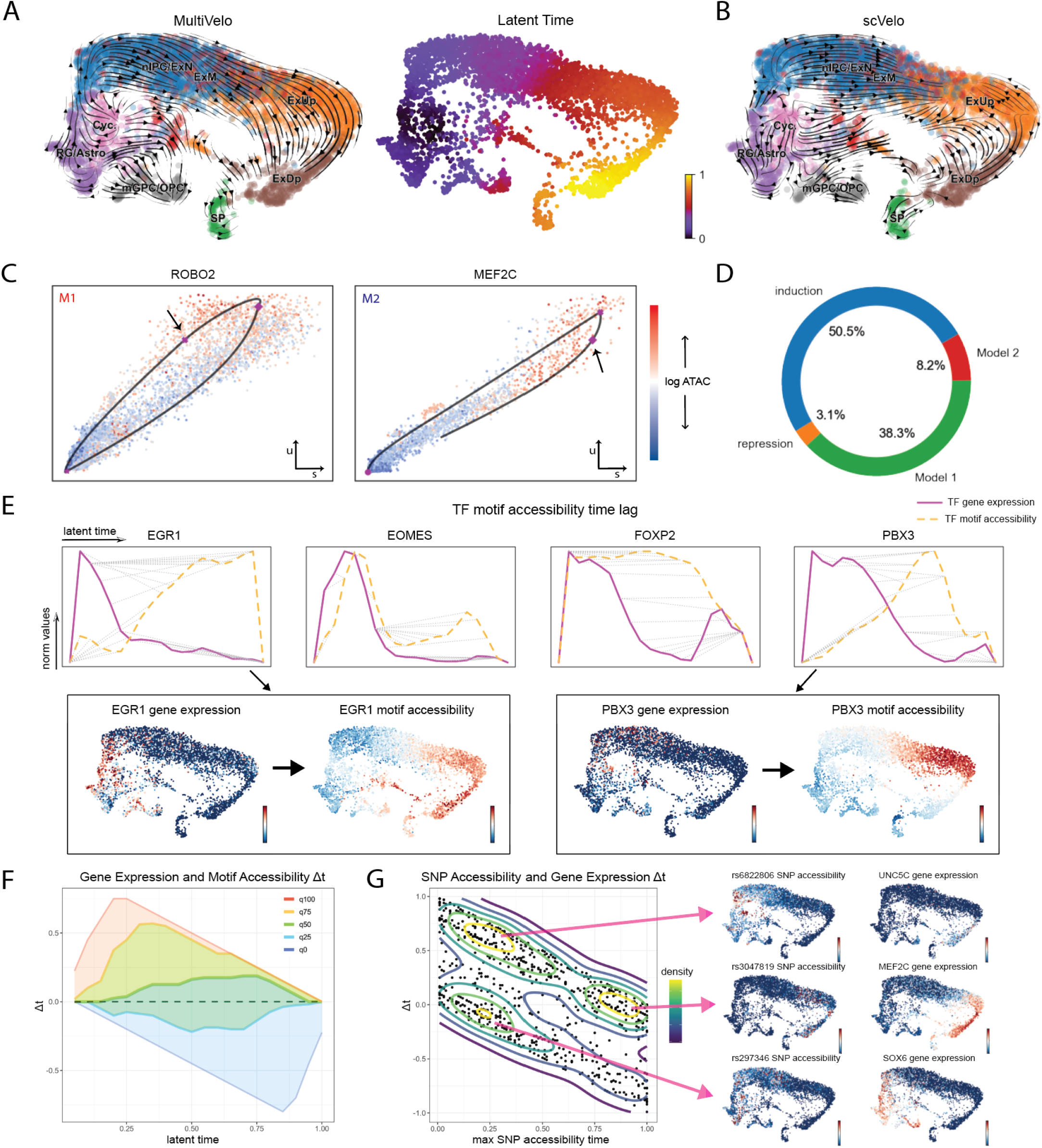
MultiVelo infers epigenome and transcriptome dynamics in embryonic human brain. **A**. UMAP coordinates with stream plot of velocity vectors (**Left**) and latent time (**Right**) from MultiVelo. **B**. Velocity streamplot from RNA-only model (scVelo). **C**. RNA phase portraits (*u* vs. *s*) colored by *c* values show clear differences between Model 1 (*ROBO2*) and Model 2 (*MEF2C*) genes. Arrows indicate where chromatin closing begins. **D**. Relative proportion of each type of kinetics across all fitted genes (n=747). **E**. Dynamic time warping alignment of TF gene expression and the accessibility of predicted binding sites for four TFs. Dotted gray lines indicate corresponding time points after alignment. Inset UMAPs colored by TF expression and motif accessibility are shown for two of the TFs, *EGR1* and *PBX3*. **F**. Quantiles of TF motif time lags inferred by DTW across all expressed TFs. The median time lag across TFs is positive at most times, indicating that TF expression generally precedes motif accessibility. **G**. Classification of SNPs according to the relationship between maximum accessibility time and time of maximum linked gene expression. The contour lines indicate density, and 3 main groups of SNPs are visible. Inset UMAP plots are shown for one example SNP from each group.

As with the mouse brain dataset, we identified clear examples of both Model 1 and Model 2 genes (Fig. 6C), though fewer genes are predicted to follow Model 2 in the human dataset (Fig. 6D). Interestingly, *MEF2C*, a Model 2 gene, is predicted by the RNA-only model to have a mostly repressive phase, likely because the “width” of the *u* − *s* phase portrait is narrow. However, the addition of chromatin information allows the correct prediction that the gene has both induction and repression phases (Fig. S6A).

A key benefit of MultiVelo is its ability to place cells onto a latent time scale inferred from both chromatin and expression data. We reasoned that latent time can identify time lags between expression and accessibility of loci other than just those immediately near a gene. For example, latent time can be used to calculate the length of time between the expression of a transcription factor (TF) and the accessibility of its binding sites (Fig. 6E and Fig. S6B-C). To do this, we used chromVar^40^ to calculate, for each cell, the total accessibility of the peaks with binding sites for each TF, subsetting to only the TFs variably expressed in the dataset. We then used dynamic time warping (DTW)^31^ to align the time series expression of each TF with the accessibility of its binding sites. This revealed a consistent pattern, in which the time of the highest RNA expression of the transcription factor preceded the time of corresponding high accessibility of downstream targets. UMAP plots colored by TF expression and binding site accessibility visually confirmed this pattern. The median time lag across all expressed TFs was positive, indicating TF expression precedes binding site accessibility in most cases (Fig. 6F). We cannot conclusively determine the mechanisms underlying these time lags without additional data. However, post-transcriptional and post-translational regulation, factors that affect the activity of chromatin remodeling complexes, and intercellular signaling could all contribute to this phenomenon.

Latent time inferred by MultiVelo is also useful for relating the chromatin accessibility of disease-related variant loci to the expression of nearby genes. We collected a list of 6968 single-nucleotide polymorphisms (SNPs) and their linked genes implicated by genome-wide association studies of psychiatric diseases, including bipolar disorder and schizophrenia. We further subset these SNPs to those overlapping chromatin accessibility peaks linked to the genes fit by our model, a total of 757 SNPs. Many of these variants occur near neuronal transcription factors and other developmentally important genes. We then calculated the chromatin accessibility, per cell, of a 400 b.p. window centered around each SNP. Using MultiVelo’s latent time, we determined the time of maximum accessibility for each SNP and the time lag between SNP accessibility and the maximum expression of its linked gene (Fig. 6G). This analysis revealed 3 major groups of SNPs, distinguished by whether their maximum accessibility occurred early or late in latent time and before or after the expression of the linked gene. UMAP plots of the SNP accessibility and linked gene expression confirm that these groups of SNPs have qualitatively distinct profiles. These groupings are significant for understanding the functions of the SNPs; for example, a SNP that is accessible only early in latent time likely plays a bigger role in developing cells than in fully differentiated cells. Similarly, a SNP whose accessibility precedes a gene’s expression is more likely to participate in regulating its expression than a SNP whose accessibility lags behind.

## 3 Discussion

In summary, MultiVelo accurately recovers cell lineages and quantifies the length of priming and decoupling intervals in which chromatin accessibility and gene expression are temporarily out of sync. Our model accurately fits single-cell multi-omic datasets from embryonic mouse brain, mouse dorsal skin, embryonic human brain, and human hematopoietic stem cells. Furthermore, our model identifies two classes of genes that differ in the relative order of chromatin closing and transcriptional repression, and we find clear examples of both mechanisms across all of the tissues we investigated. We anticipate that MultiVelo will provide insights into epigenomic regulation of gene expression across a range of biological settings, including normal cell differentiation, reprogramming, and disease.

## 4 Methods

### 4.1 Previous Approaches: RNA velocity

In the original RNA velocity model, the proposed system of differential equations for RNA splicing is as follows

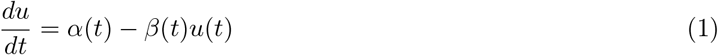

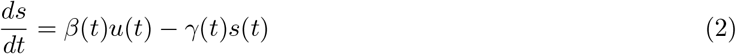

where *u* is unspliced RNA, *s* is spliced RNA, and *α, β, γ* are transcription, splicing, and degradation rate respectively. Assuming constant transcription and degradation rates, the rate equation parameters can be normalized by *β* and are reduced to

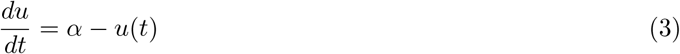

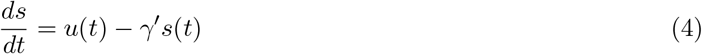

In steady-state cell populations, the amount of spliced mRNA does not change: 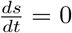. Therefore, 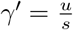 and *α* = *u*. The ratio *γ*^*′*^ can be calculated using a simple linear regression that fits cells with expression values in upper and lower quantiles. RNA velocity is then defined as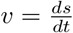.

Bergen et al. developed a dynamical RNA velocity model (scVelo) by extending the original equations to include time and cell state latent variables, capturing transient states between steady states.

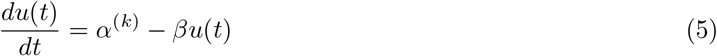

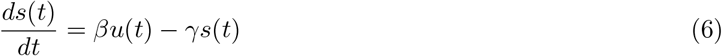

where *k* indicates one of the four transcription states: induction (*k* = 1), repression (*k* = 0), and two associated steady states (*k* = *ss*1 and *k* = *ss*0).

This system of differential equations can be solved analytically as follows:

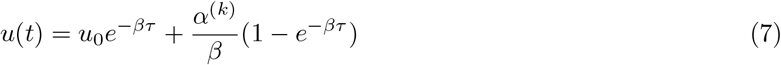

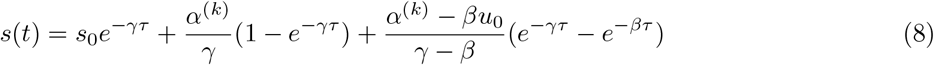

where *u*_0_ and *s*_0_ are initial values, and 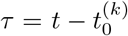 is the time interval from the start of the induction or repression state.

The analytical solution converges to the steady-state values as *τ* →− ∞:

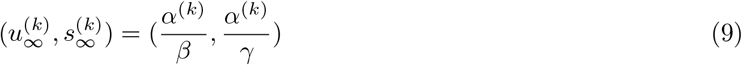

Because the equations involve the latent time variable *τ*, scVelo uses an expectation maximization algorithm to iteratively estimate latent time and the parameters of the ODE *θ* = (*α*^(*k*)^, *β, γ*), as well as state starting time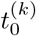. Cells are assigned to latent times by approximately inverting the ODE solution.

### 4.2 Differential Equation Model of Gene Expression Incorporating Chromatin Accessibility

To incorporate chromatin accessibility measurements into a differential equation model of gene expression, we assume that the rate of transcription for a gene is influenced by the accessibility of its promoter and enhancers. For simplicity, we model a single value *c*, which is the sum of accessibility at the promoter and linked peaks for a gene. Unlike gene expression, which can theoretically grow without bound, it is possible in principle for chromatin to be fully open or fully closed at a particular locus. Thus, we normalize chromatin accessibility to [0, 1], and assume that *c* approaches 1 with rate of change proportional to *α*_*co*_ *>* 0 during the opening phase and approaches 0 with rate of change proportional to *α*_*cc*_ *>* 0 during the closing phase. Our biological motivation for this mathematical formulation can be summarized as follows: impulses of remodeling signals cause chromatin to begin opening or closing rapidly at first. However, biochemical constraints such as the structures of histone complexes and their inter-molecular interactions gradually slow the rate of opening or closing so that *c* asymptotically approaches full accessibility or inaccessibility (Fig. S3A). Empirically, we find that the observed *c*(*t*) values in single-cell multi-omic dataset show this qualitative behavior (Fig. S3B). We define a new system of differential equations to reflect these modeling assumptions:

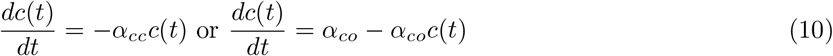

If we assume that the chromatin opening and closing kinetics are mirror images of each other, only a single chromatin rate parameter *α*_*c*_ *>* 0 is required, and the system of equations simplifies to:

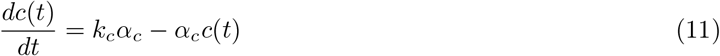

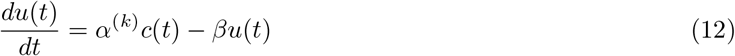

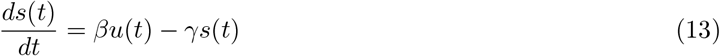

where

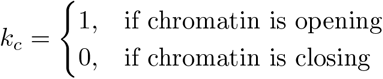

As with the RNA velocity model, we define chromatin velocity as 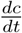. The parameter *k*_*c*_ allows for different dynamics during chromatin opening (*k* = 1) and chromatin closing (*k* = 0), analogous to how the transcription rate *α*_*k*_ in the dynamical RNA velocity model varies between transcriptional induction and repression phases (*k* = 1 and *k* = 0). The system of differential equations can be solved analytically to obtain:

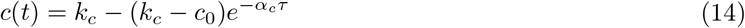

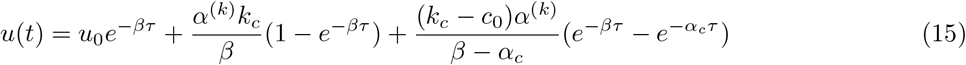

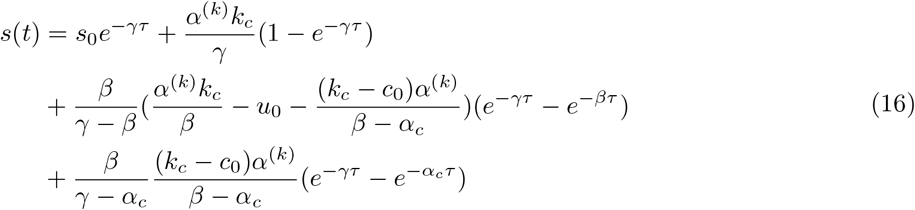

where *c*_0_, *u*_0_, and *s*_0_ are the initial values of one of the four states, and *τ* = *t* − *t*_0_ is the time interval from the start of that state. Note that the analytical solution is the same even if we assume different opening and closing rates, if we simply use

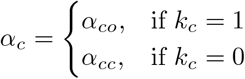

Similar to RNA velocity, the origin of the trajectory is (0, 0, 0) (whether observed or not), and initial values of the next state can be obtained by solving the expected values at the switch interval using equations for the previous state. The range of chromatin values is restricted to [0,1] to span from fully closed to fully open chromatin accessibility. As such, the hypothetical steady states for chromatin accessibility 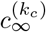, as time approaches infinity on each interval, is simply 0 for closing state and 1 for opening state. The steady-state values for each state become

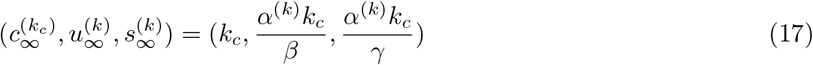

Because the model includes separate latent variables for chromatin state *k*_*c*_ and RNA state *k*, there are multiple potential orders of chromatin remodeling states and transcription states. We label these possible orders as Model 0 (M0), Model 1 (M1), and Model 2 (M2):

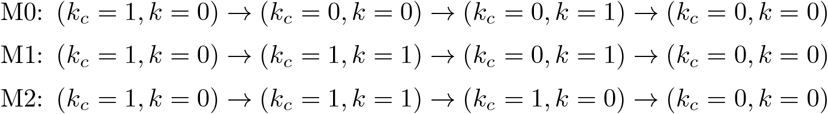

We reason that it is biologically implausible for chromatin to be closed when transcription initiates, because it is difficult or impossible for a gene with inaccessible chromatin to be transcribed. Thus, we implement the capability to fit Model 0 if desired, but fit only Model 1 and Model 2 by default. Model 1 and Model 2 are both biologically plausible, and these different orders have biologically meaningful interpretations. We refer to Model 1 as delayed transcriptional repression and Model 2 as delayed chromatin repression. Within each model, a trajectory is defined by a set of eight core parameters *θ*, including three phase switching time points (transcriptional initiation time *t*_*i*_, chromatin closing time *t*_*c*_, and transcriptional repression time *t*_*r*_) and five rate parameters (chromatin opening rate *α*_*co*_, chromatin closing rate *α*_*cc*_, transcription rate *α*, splicing rate *β*, and RNA degradation rate *γ*). There is also a fourth possible switch time *t*_*o*_ at which chromatin opening begins, but by excluding Model 0 we can assume that *t*_*o*_ = 0 for all genes.

### 4.3 Model Likelihood

We can formulate a probabilistic model to calculate the likelihood of the observed data for a gene under particular ODE parameters *θ*. To do this, we simply assume that the observations are independent and identically distributed, and that the residuals are also normally distributed with mean given by the deterministic ODE solution and diagonal covariance. Because we scale the *c, u*, and *s* values, we can further assume that the variance is the same in all directions. That is, if we define the ODE prediction as 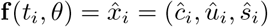, then the distribution of the observed data **x**_**i**_ = (*c*_*i*_, *u*_*i*_, *s*_*i*_) for each gene is:

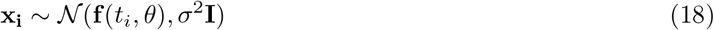

The negative log likelihood of all *n* observations is then

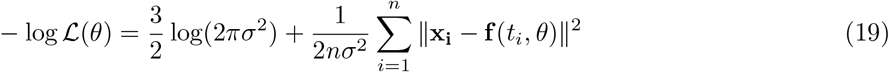

We can infer the ODE parameters *θ* by maximum likelihood estimation, which is equivalent to minimizing the mean-squared error. The maximum likelihood estimate of *σ*^2^ is the sample variance of the residuals along each coordinate. We can then rank genes by their likelihood to identify the genes best fit by the ODE model. We can also determine which model best explains the *c, u, s* values observed for a particular gene by comparing the mean squared error (MSE) under Model 1 and Model 2.

### 4.4 Parameter Estimation and Latent Time Inference by Expectation Maximization

Both the cell times *t* and the ODE parameters are unknown, so we perform expectation-maximization to simultaneously infer them. The E-step involves determining the expected value of latent time for each cell given the current best estimate of the ODE parameters. Because inverting the three-dimensional ODEs analytically is not straightforward, we perform this time estimation by finding the time whose ODE prediction is nearest each data point, selecting the time from a vector of uniformly spaced time points (see Implementation Detail section). In the M-step, we find the ODE parameters that maximize the data likelihood (equivalent to minimizing MSE) given the current time estimates for each cell. We use the Nelder-Mead simplex algorithm to minimize MSE.

### 4.5 Model Pre-Determination and Distinguishing Genes with Partial and Complete Dynamics

A gene does not have to complete a full trajectory within the measured cell population. In fact, for differentiating cells, we found that it is not uncommon for a gene to possess only an induction or repression phase, especially for differentially expressed cell-type marker genes. The three types of gene expression patterns (induction only, repression only, and complete trajectory) can be directly inferred before fitting a model, thus avoiding ambiguous assignments near RNA phase transition points.

We used a combination of two methods for this purpose. The first method directly results from the assumptions of RNA velocity: given a steady-state fit, cells in the induction phase reside above the fitted steady-state line while cells in the repression phase reside below the steady-state line. Thus, the ratio of sum of squared distances (SSE) of cells on either side of the steady-state line is an indicator that can be used to determine the direction of the trajectory.

The second method incorporates low-dimensional coordinates (e.g., from PCA or UMAP) as global information. We use UMAP coordinates by default, because these are often precomputed for visualization. Assuming that a gene possesses a complete trajectory, then at lower quantiles of its unspliced-spliced phase portrait, these cells are expected to have a bimodal pairwise distance pattern in the low-dimensional representation. Such a bimodal pattern indicates dissimilar populations, as some of these cells are in the early phase of induction, while the others have reached the late phase of repression. In contrast, for partial trajectories, cells at lower quantiles of the RNA phase portrait will have similar low-dimensional coordinates. Similarly, the unimodal or bimodal pattern can also be derived from the assumption that noise is normally distributed along the trajectory given by the ODE solution. We thus used a Gaussian mixture model to test if the distribution of pairwise distances among cells in a gene’s lower quantile region is unimodal or bimodal, designating the trajectory being partial or complete, respectively. In order to be classified as a complete trajectory, the distance of the means between two Gaussians under bimodal distribution must exceed the globally measured variation (one standard deviation by default) of all pair-wise distances on the low-dimensional coordinates for cells that express that gene, and the weight of the second, usually smaller Gaussian must pass a certain threshold (0.2 by default). The final assignment of partial or complete trajectory utilizes a combination of both methods (steady-state line ratio and bimodality), with the first method given priority.

Additionally, whether a gene is better explained by Model 1 or Model 2 can be determined without actually fitting parameters under both models. To see how, note that the chromatin closing phase precedes transcriptional repression in Model 1 but succeeds transcriptional repression in Model 2. This implies that the highest chromatin accessibility values occur during the transcriptional induction phase for Model 1 genes but during the repression phase for Model 2 genes. Thus, the ratio of top chromatin values across the steady-state line can be used to determine whether each gene is best described by Model 1 or Model 2 before actually fitting the parameters. We implement this model pre-determination as a default to speed up computation, but users can alternatively opt to fit both models and compare their losses instead.

### 4.6 Parameter Initialization

Parameters specifically related to RNA (*α, β, γ*, and the RNA switch time interval) are initialized based on steady-state model as in scVelo. The rescaling factor for chromatin accessibility is initialized to 1, as the maximum observed accessibility is likely some value in-between 0 and 1. Other parameters can be found in Implementation Detail section below.

We also initialize a scale factor for *u*. Here we show that its value is closely related to the roundness of the U-S portrait under steady-state assumptions. First, *u* and *s* are both normalized to the range [0, 1]. Next, points of steady-state rate are found on the induction phase

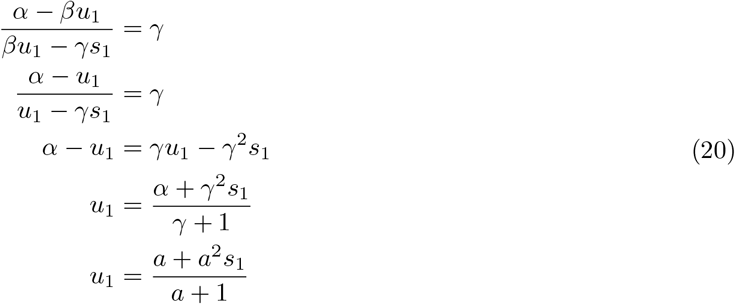

where *a* is an unknown scalar and equals to the expected maximum of rescaled *u*. And similarly on the repression phase

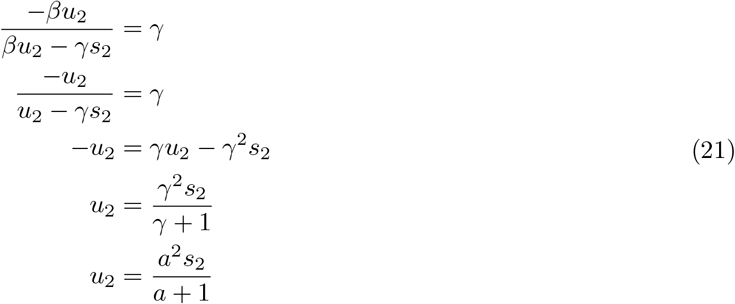

Then if we assume 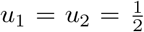 of maximum unspliced count, meaning the line connecting *u*_1_ and *u*_2_ is parallel to *s*-axis and at the same time, crosses the middle point of *u* (due to symmetry), then:

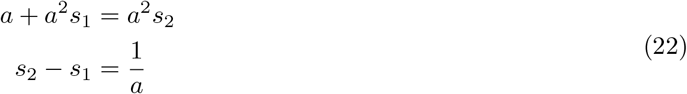

The rescale factor for u is therefore *s*_2_ − *s*_1_ around middle of u when s is normalized to range of [0, 1]. *u/*(1*/a*) = *a* ∗ *u* and *s* are then used to initialize other parameters. Note that value of *a* is then further optimized during fitting.

### 4.7 Implementation Detail

A key implementation detail is how to estimate each cell’s latent time given the ODE solution from the current parameters. Inverting the ODE solution is analytically challenging due to the complexity arising from a system of 3 ODEs. Thus, rather than pursuing an exact or approximate analytical solution to calculate time, we simply maintain a set of anchor points uniformly spaced in time. For each cell, we then identify the nearest anchor point and assign the cell’s time to the time of the anchor point. In more detail, we calculate the (*c, u, s*) values of the ODE solution at a specified number of uniformly distributed time points. Then we calculate pairwise distances from the observed cells to these anchor points. The shortest distance represents the residuals to the inferred trajectory, and the time of the anchor point is assigned to the cell. We found that 500-1000 points are sufficient to capture the full trajectory dynamics. We restrict the time range to span from 0 to 20 hrs, consistent with scVelo’s default setting.

After determining trajectory direction and model to fit, expression values are shifted so that the minimum value starts from zero, then they are scaled but not centered. RNA rate parameters are initialized based on the steady-state model: *α* is initialized as the mean of top-percentile *u* values to represent a gene’s overall transcription potential^7^. The splicing rate *β* is initialized to 1–consistent with the steady-state model heuristic–and the degradation rate *γ* is obtained through linear regression of the top-percentile (*u, s*) values^6^. Chromatin rate *α*_*c*_ is initialized as − *log*(1 − *c*_*high*_)*t*_*sw*3_ where *c*_*high*_ is the mean accessibility of those cells with accessibility above average of all cells for that gene, and *t*_*sw*3_ is the chromatin closing switch time in the current grid search iteration. We initialize the RNA switch-off time using the explicit time-inversion procedure described in scVelo’s method. To initialize the RNA switch-on time and chromatin switch-off time, we search over a grid of times 2 hrs apart. The best initial switch time combinations are chosen based on mean squared error loss.

To fit and optimize parameters, we minimize the negative log likelihood (equivalent to MSE loss) using the Nelder-Mead downhill simplex method^41^, implemented in the scipy minimize function. The Nelder-Mead algorithm performs a series of transformations on the model parameters, including reflection, shrinking, and expansion to improve the fitting results. When fitting induction-only trajectories, only the first two phases (chromatin priming phase and coupled induction phase) are aligned to observations. When fitting repression-only trajectories, only the later two phases are fitted. To improve convergence speed, we minimize with respect to subsets of parameters at any time, holding the others fixed. This is similar to a block coordinate descent strategy. Within each iteration, we first update parameters exclusive to *c*, then parameters related to *u*, and finally parameters affecting *s*. We found that 5-10 iterations are sufficient for convergence in most cases. To ensure that the switch times occur in the proper order (e.g., transcriptional induction precedes transcriptional repression), we opted to use switch intervals rather than switch time-points as actual parameters. Thus a model is guaranteed to be valid if all parameters are positive, with no other constraints needed.

The trajectory constructed using a set of rate parameters is represented by a set of uniformly distributed anchor time-points. By using the uniform distribution, we assume cells have equal prior probability to be measured at any given time-point. The local sparsity of cells is determined by model parameters. We used KD-tree^42^ from scipy to search for the closest anchor to each observation and its corresponding distance. Using anchor points also allows the model to mimic the expected local sparsity of cells along the fitted trajectories by encouraging anchors to concentrate near where cells concentrate in order to reduce small distance offsets caused by discrete representation of the trajectory.

After fitting the models, because genes with partial fitted trajectories result in a shorter total observed time-range–violating the assumption that all genes share one time scale–the rate parameter set and the switch times are scaled down and up, respectively, so that time ranges from 0 to 20 hr. (Note that multiplying the time and dividing the rates by the same constant will result in identical trajectories.) This ensures that the time parameters from all genes are comparable. Switch times are shifted backward in time if the observable start of the trajectory happens later than 0 hr.

The optimized rate parameters and time assignments are plugged back into the system of ODEs to obtain velocities for chromatin accessibility, unspliced RNA, and spliced RNA for each cell. Our multi-omic velocity method is implemented in python. Many internal functions in our method have been accelerated with Numba. Distances, time assignments, and velocity vectors are smoothed among nearest neighbors to mitigate the effect of measurement stochasticity.

Because multi-omic velocity is an upstream extension of the original RNA velocity model, it can be easily reduced to the RNA-only model by setting chromatin to be fully open (constant of 1) throughout the entire trajectory. Fitting this RNA-only model is then very similar to running the multi-omic model, but there will be no notion of the Model 1 and Model 2 distinction.

### 4.8 Post-fitting Analyses

Bergen et al.^7^ have developed great downstream analyses methods for RNA velocity in the scVelo toolkit. Because our method is a direct extension of the dynamical model to multi-omic data, many of scVelo’s methods can be applied with only a change of arguments. Our main method replaces the scVelo functions tl.recover_dynamics and tl.velocity. In this paper, scVelo’s tl.velocity_graph with total-normalized spliced velocity vectors computed from our multi-omic method was used to obtain a transition matrix between cells based on cosine similarity between a cell’s velocity vector and expression differences. We used pl.velocity_embedding_stream to embed and plot velocity streams onto UMAP coordinates. Computation of global latent time among cells and genes is implemented in tl.latent_time.

We performed Dynamic Time Warping using the dtw R package^43,44^. First, the accessibilities or expressions of cells were aggregated to 20 equal-sized bins based on either their gene time (for *Wnt3* in the skin dataset) or latent time (for human brain motifs), and then maximum-normalized to the same range of [0, 1]. For motifs, a rolling mean of three-bin was applied to the RNA and motif counts to smooth the curves. We then added a zero to each end of the time series to ensure that the starting and ending values of each time series matched. Then we used dtw to find the best alignment–local for *Wnt3* or global for motifs–between the two time series with Euclidean distance penalty. We then calculated time lags by simply subtracting the times of the aligned points. When many-to-one mappings occurred in global alignments, we averaged the time lags across all points mapped to the same time. For SNP time analysis, both the SNP accessibilities and log RNA expressions were aggregated to 100 equal-sized bins. We then calculated the time lag as the time difference between the time bins with highest values in the two modalities.

### 4.9 Generation of Simulated Data

1000 genes were simulated with various rate parameters, switch times, time sequences, and models (1 and 2). *α*_*c*_, *α, β*, and *γ* values were generated from multivariate log-normal distributions with mean −2, 2, 0, 0 and variance 0.5, 1, 0.3, and 0.3, with a small covariance of 0.01 between *α*_*c*_, *α* and *β*. Four switch intervals were random chosen from [1,4], [1,9], [1,9], and [1,9], and scaled to give a time range from 0-20 hrs. The model (Model 1 vs. Model 2) was sampled uniformly at random. Cell times were sampled from a Poisson distribution. Noise was added to each cell with diagonal covariances of [*max*(*c*)^2^*/*90, *max*(*u*)^2^*/*90, *max*(*s*)^2^*/*90]. The accuracy of loss-based and predetermined model decisions were separately computed.

### 4.10 Preprocessing of data, weighted nearest neighbors, and smoothing

#### 10X embryonic E18 mouse brain

Filtered expression matrix for ATAC-seq, feature linkage file, as well as position-sorted RNA alignment (BAM) file of E18 mouse embryonic brain data of around 5k cells were downloaded from 10X Genomics website (CellRanger ARC 1.0.0). Total, unspliced and spliced RNA reads were separately quantified using the Velocyto run10x command. The resulting loom file was read into python as an AnnData object and preprocessed with scanpy and scVelo to perform filtering, normalization, and nearest neighbor assignment. Next, clusters were computed using the Leiden^45^ algorithm. Cell-types were manually annotated based on expression of known marker genes^46,47,48,49^. We then excluded interneurons, Cajal-Retzius, and microglia cell populations for our downstream analyses, because these cell types are not actively differentiating. We then re-processed the raw counts of subset clusters, which consists of more than 3k remaining cells, with scVelo. The unspliced and spliced reads were neighborhood smoothed (averaged) by scVelo’s pp.moments method with 30 principal components among 50 neighbors. The downloaded feature linkage file contains correlation information for gene-peak pairs of genomic features across cells. We first collected all distal putative enhancer peaks (not in promoter or gene body regions) with ≥ 0.5 correlation with either promoter accessibility or gene expression that were annotated to the same gene or within 10kb of that gene. We then aggregated these enhancer peaks with 10X annotated promoter peaks for the corresponding genes, as a single chromatin accessibility modality to boost chromatin signal. These aggregated accessibility values were then normalized using the term frequency–inverse document frequency (TF-IDF) method^24^. (Note that during fitting, chromatin values are normalized to [0, 1], so using other total-count based normalization will produce identical results.) Due to the increased sparsity of ATAC-seq data, the neighborhood graph and clustering results based solely on peaks is often noisy and unreliable. Seurat group recently developed a method to compute neighborhood assignments for simultaneously measured multi-modality data in the Seurat V4 toolkit, which they called weighted nearest neighbor (WNN)^50^. The WNN method learns weights of each cell in either modality based on its predictive power by neighboring cells in each of the modalities, so that both RNA and ATAC information can be incorporated when assigning neighbors. We used 50 WNNs obtained from Seurat for each cell to smooth the aggregated and normalized chromatin peak values. Our WNN analysis followed the recommended steps in Seurat V4 vignette for 10X RNA + ATAC. We thus obtained three matrices containing chromatin accessibility, unspliced, and spliced counts. Shared cell barcodes and genes were filtered among matrices and resulted in 3365 cells and 936 highly variable genes, these matrices were then used for dynamical modeling.

#### SHARE-seq mouse skin (hair follicle) data

The quantified ATAC-seq expression matrix, raw ATAC-seq fragments file, and cell annotations of SHARE-seq mouse skin dataset^9^ were downloaded from GEO: GSE140203. The RNA alignment BAM file as well as UMAP coordinates for TAC, IRS, Medulla, and Hair Shaft Cuticle/Cortex cell populations used in the SHARE-seq manuscript were obtained directly from the authors. We run Velocyto to quantify unspliced and spliced counts, and the RNA AnnData object was further preprocessed with scanpy/scVelo for the four cell types of interest. In R, the chromatin fragment file was used to construct a gene activity matrix by aggregating peaks onto gene coordinates using the GeneActivity function in Signac. Domain of regulatory chromatin (DORCs) is defined as chromatin regions that contain clusters of peaks that are highly correlated with gene expressions in SHARE-seq’s analysis. A list of computed DORCs coordinates was downloaded from its supplementary material section. These coordinates were output to the bed format, and we extracted fragments together with their corresponding cell barcodes that overlap with these DORCs regions. A peak expression matrix for DORCs was constructed with Liger’s makeFeatureMatrix method. The gene activity and DORCs counts were then merged in python to form a single chromatin modality. Similar to brain data, this matrix underwent TF-IDF normalization and WNN smoothing. A total of 6436 cells and 962 genes participated in the downstream analyses.

#### Human hematopoietic stem and progenitor cell (HSPC)

Purified human CD34^+^ cells were purchased from the Fred Hutch Hematology Core B. Freshly thawed cells were maintained at 37°C with 5% CO_2_ in Stemspan II medium supplemented with 100 ng/ml stem cell factor, 100 ng/ml thrombopoietin, 100 ng/ml Flt3 ligand (all from Stemcell Technologies), and 100 ng/ml insulin-like growth factor binding protein 2 (R&D Systems) for seven days. HSPCs were prepared according to the manufacturer’s “10X Genomics Nuclei Isolation Single Cell multiome ATAC + Gene Expression Sequencing” demonstrated protocol. Briefly, cells were washed in PBS supplemented with 0.04% BSA and sorted using the Sony SH800 cell sorter (Sony Biotechnologies). Nuclei were isolated following the “Low Cell Input Nuclei Isolation” sub-protocol and immediately processed using the Chromium Next GEM Single Cell Multiome + Gene Expression kit.

10X filtered expression matrices, Velocyto computed unspliced and spliced counts, and feature linkage and peak annotation files from CellRanger ARC 2.0.0 were read into python to construct RNA and ATAC AnnData objects. Filtering, normalization, and variable-gene selection were performed following scVelo’s online tutorial. Because HSPCs are rapidly proliferating, we noticed systematic differences in cell cycle stage across the set of cells. The cell-cycle scores for both G2M and S phases, computed using scVelo’s tl.score_genes_cell_cycle function were then regressed out of the RNA expression matrices with scanpy’s pp.regress_out function (Fig. S5B). Note that the regression did not change unspliced and spliced counts. Then gene expression scaling was performed. ATAC peaks were aggregated and normalized using the same procedure as described for the 10X mouse brain. Joint filtering between RNA and ATAC resulted in 11605 cells and 1000 genes. RNA expression was smoothed by scVelo’s pp.moments with 30 principle components and 50 neighbors. Leiden found 11 clusters. Cell types were assigned based on canonical HSPC markers^51,52,53,54,55^. The chromatin accessibility matrix was WNN smoothed with 50 neighbors computed using Seurat. Then the RNA and ATAC objects were input to our dynamical function with default parameters. We relaxed the likelihood threshold for velocity genes (used for computing the velocity graph) to 0.02 compared to the default of 0.05 due to noisiness of this dataset.

To find complete genes in each of the lineages from HSC towards GMP (myeloid), erythrocytes, and platelets, we subset cells of each specific lineage and select known complete genes as those genes that have higher unspliced and spliced expressions in the progenitor populations leading to each of the terminal cell types. We then ran the model predetermination algorithm based on peak chromatin accessibility as described in the previous section. The genes predicted as Model 1 and Model 2 for each lineage are then merged with duplicates removed, and we performed gene ontology enrichment analysis (GOrilla^56^) using all sequenced genes as the background set.

Preprocessed bulk ChIP-seq peaks of H3K4me3, H3K4me1, and H3K27ac for CD34+ HSPC were downloaded from GSE70677^37^. Peaks were mapped to genes with Homer^57^. Known complete genes in the myeloid and erythroid lineages were grouped together, and predicted M1 and M2 genes were extracted. Scores of peaks associated with the same genes were aggregated. Wilcoxon rank-sum test was used to compute significance.

#### Human cerebral cortex

We obtained the multiome RNA, unspliced, spliced, and ATAC-seq peak files from the authors. The ATAC peak matrix contains consensus peaks of non-overlapping uniform 500bp length. After initial clustering, we observed a severe batch effect in one of the three samples. We thus decided to removed this third sample and perform all downstream analyses with the two remaining samples (dc2r2_r1 and dc2r2_r2). We re-named the clusters from the original paper as follows based on marker gene expression: RG → RG/Astro, nIPC/GluN1 → nIPC/ExN, GluN3 → ExM, GluN2 → ExUp, GluN4 and GluN5 → ExDp^47^. Peaks were annotated to genes with Homer^57^. We considered peaks within 10000bp of transcription start sites as promoter peaks. A list of peak-gene links and correlations were downloaded from the supplementary material and aggregated to promoter peaks if the correlation exceeded 0.4. After filtering the RNA and ATAC matrices, 4693 cells and 919 genes were left and input to model fitting. TF motif profiles were computed with chromVAR^40^ on the JASPAR2020 database^58^ using all consensus peaks. The background-corrected deviation z-scores were used as normalized motif accessibilities, and the values were smoothed with WNN. Then TF genes appearing in the variable gene list (after internal filtering by the dynamical function) were extracted for time-lag analysis, which resulted in 30 known motifs. All mental or behavioural disorder associated SNPs (EFO_0000677) were downloaded from the Ensembl GWAS Catalog. The list contains 6968 SNPs, and filtering for overlap with consensus peaks linked to the top genes resulted in 757 SNPs. Each SNP’s accessibility was quantified as the count of all ATAC fragments that overlap a 400 b.p. bin centered on the SNP location. The accessibility matrix was normalized by library size and smoothed by WNN neighbors.

## 5 Code and Data Availability

MultiVelo is implemented in Python. The package is available on GitHub (https://github.com/welch-lab/MultiVelo) and PyPI. The newly sequenced 10X Multiome HSPC sample will also be uploaded to dbGAP and GEO.

## 6 Acknowledgements

This work was supported by NIH grants R01AI149669 to KLC and JDW and R01HG010883 to JDW. The HSPC sample was provided by Cooperative Center of Excellence in Hematology grant DK106829. We thank Jun Li, Stephen CJ Parker, Yichen Gu, and members of the Collins lab for helpful discussions.

