## Supplemental Figures for "Single-cell multi-omic velocity infers dynamic and decoupled gene regulation"

### 7 Supplementary Material

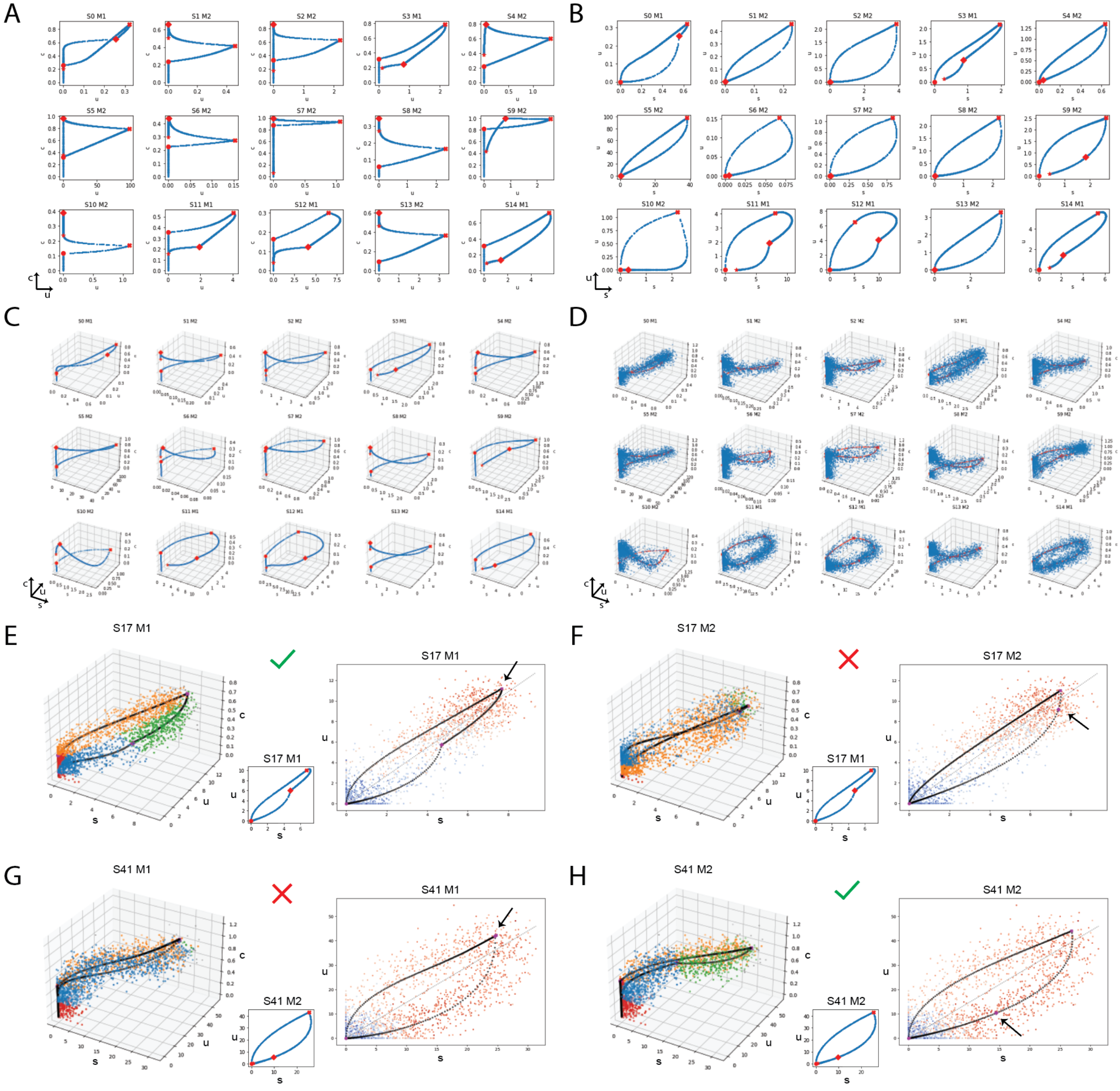


**Fig. S1. Simulated Data Analysis.** A total of 1000 genes were simulated with various parameters for both model 1 and model 2. **A**. C-U view of noiseless simulations of 2000 time-points in the 0-20 hr range. **B**. U-S view of noiseless simulations from A. **C**. 3D view of noiseless simulations from A. **D**. Noise added to simulated points to mimic real data. **E** and **F**. Model 1 and Model 2 fits for the same simulated gene (S17). The likelihood is higher under Model 1, consistent with the ground truth. **E**. **Left**: 3D view of the fitted Model 1 trajectory colored by states, along with predicted switch time points. **Middle**: simulation with ground-truth switch times. **Right**: U-S view of fitted trajectory colored by $log(c)$. **F**. Similar to **E**, but the fit shown is for Model 2 (the incorrect model). **G** and **F**. Model fits for simulated gene S41, similar to **E** and **F**, but this time, Model 2 is the ground truth model. MultiVelo correctly identifies the sample to be Model 2 with accurate switch time estimations. The model assignments of 985/1000 samples were correctly predicted based on likelihood.

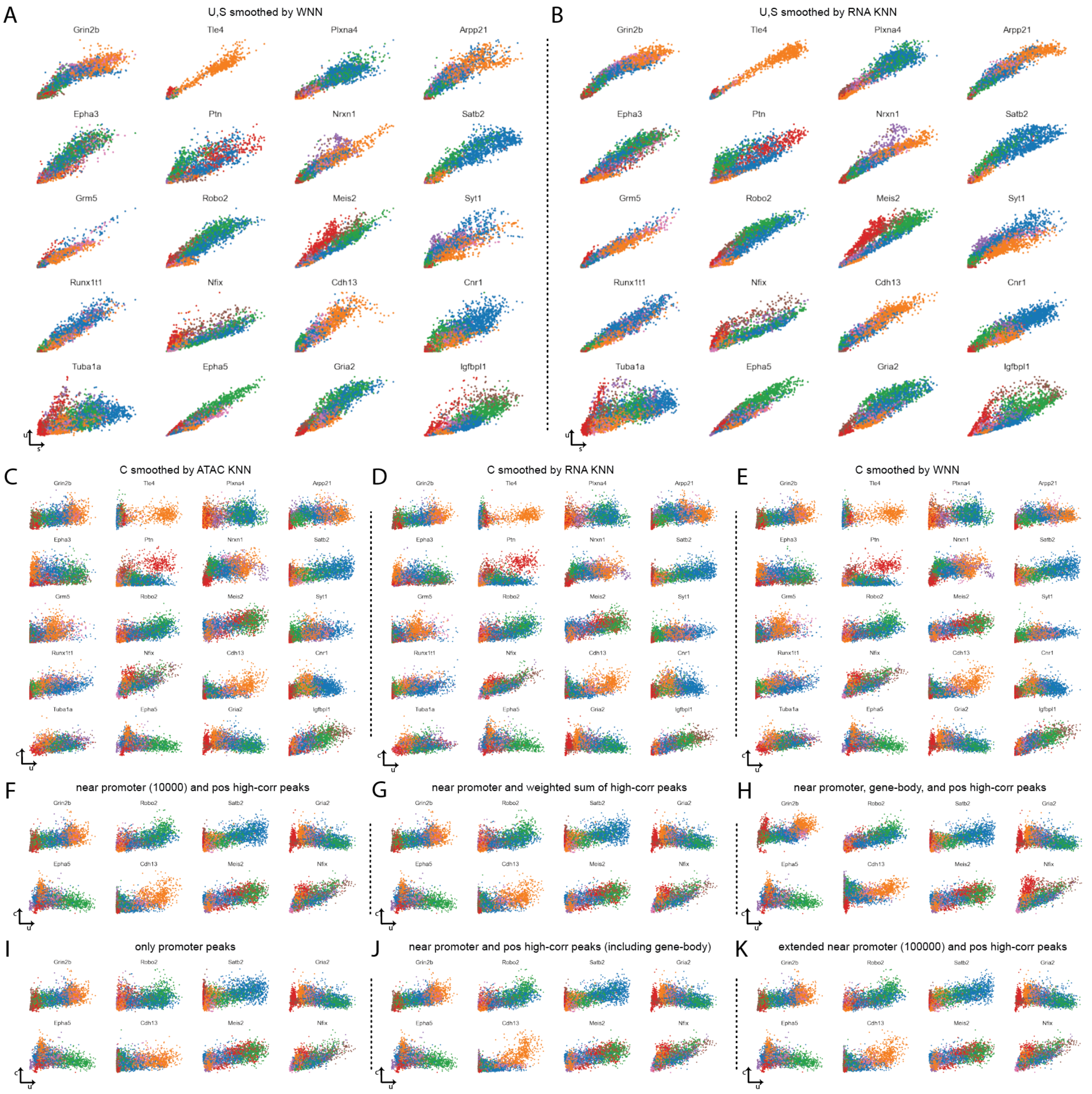


**Fig. S2. Comparison of different KNN smoothing and ATAC-seq aggregation methods. A**. RNA smoothed by weighted nearest neighbors from Seurat. **B**. RNA smoothed by RNA neighbors–the same as scVelo (current chosen method). **C**. TF-IDF normalized ATAC-seq counts smoothed by neighbors in latent semantic indexing (LSI) space. **D**. Similar to **C** but with RNA k nearest neighbors. **E**. Similar to **C** but with weighted nearest neighbors. WNN approach is chosen to be the current method because it provides a balance between RNA and ATAC information. **F**-**K**. Comparison of strategies for aggregating peaks to a single $c$ value per gene. **F**. All peaks near promoter (10000 bp) annotated to the target gene and positively correlated peaks with either promoter peaks or gene expression are added. **G**. Similar to **F** but the addition of peaks is weighted by their correlation. Negatively correlated peaks are also weighted summed with negative correlations **H**. Similar to **F** but also include peaks in the target gene's body. **I**. Only included promoter peaks. **J**. Similar to **F** but also include highly correlated peaks that are within or near other genes. **K**. Similar to **G** but with extended range of 100000 bp.

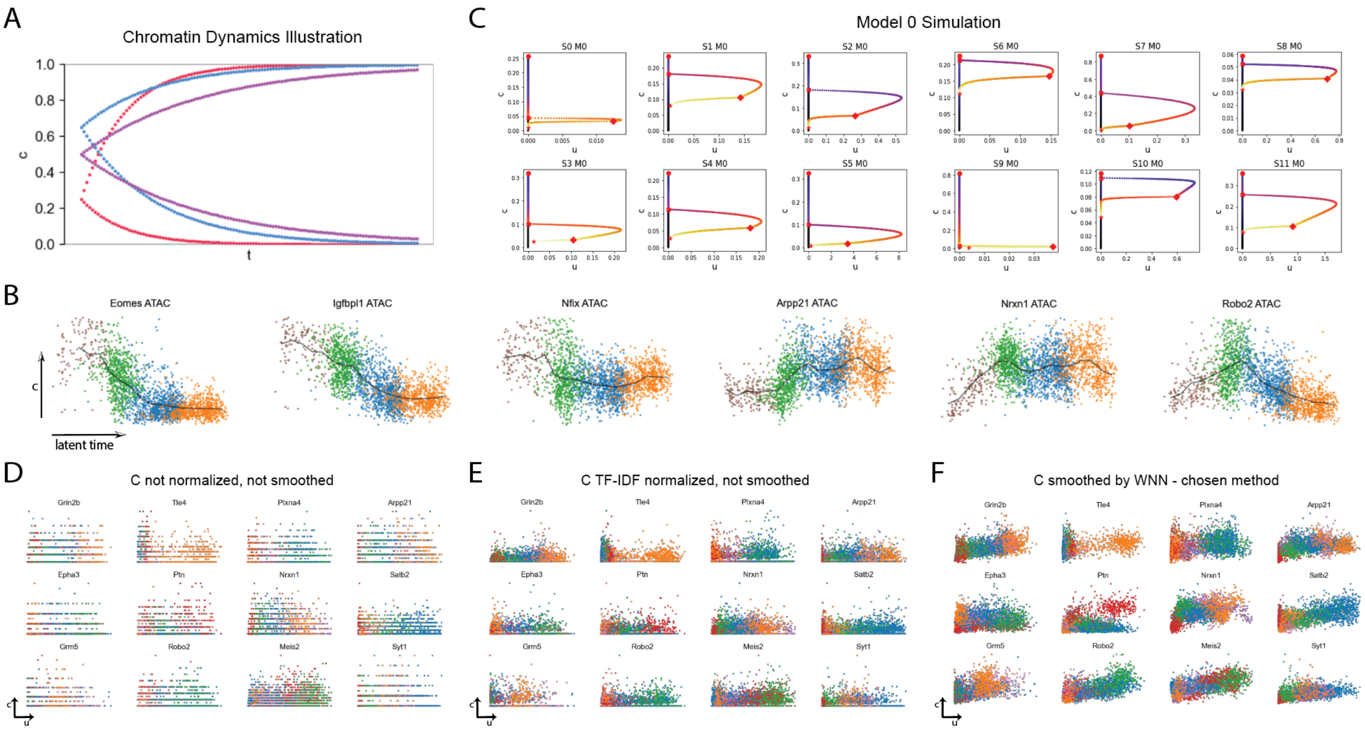


**Fig. S3. Chromatin dynamics, Model 0, and necessity of chromatin data preprocessing. A**. Chromatin dynamics illustration: chromatin opening and closing are modeled as asymptotically approaching fully opened (1) or fully closed (0) starting from any initial value. **B**. Chromatin accessibility change as a function latent time inferred by scVelo using only the RNA portion of the 10X multiome mouse brain dataset (colored by mouse brain cell types). Black lines connect the mean accessibilities within 20 equal-sized windows. The shapes of the ATAC trends are qualitatively very similar to the ODE model we propose. **C**. Simulation of Model 0 samples. The long delay between chromatin closing and transcription initiation is unlikely to happen in real biological systems. In the rare cases when high chromatin accessibility but low expression or high expression but low accessibility pattern is observed, it is likely due to technical issues such as dropout or background noise. **D**. The need for normalization as a preprocessing step for ATAC-seq. **E**. The need for smoothing as a preprocessing step for ATAC-seq. **F**. Chromatin accessibility results after peak-to-gene aggregation, TF-IDF normalization, and WNN smoothing. It is the same as Fig. S2E.

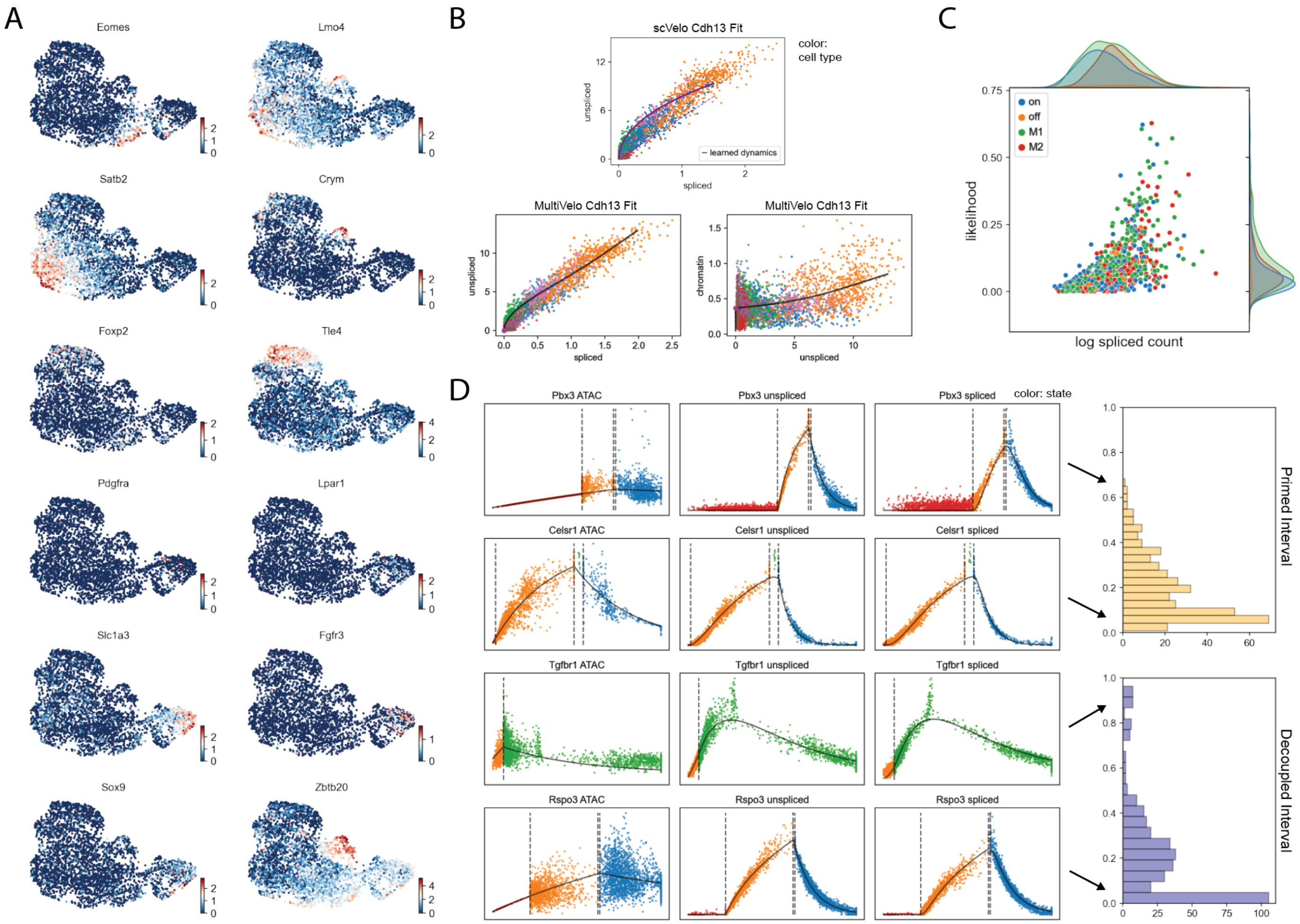


**Fig. S4. Additional figures for mouse brain. A**. Canonical marker gene expression for embryonic mouse brain cell types. **B**. Comparison of Cdh13 fits from scVelo and MultiVelo. An elevating transcription rate due to opening of chromatin produces a more linear fit and better captures the observed phase portrait. **C**. Scatterplot of gene likelihood against log total spliced count. Gene likelihood is not significantly affected by model assignment or trajectory type. Likelihood does increase with spliced count, as this usually indicates higher quality or highly-variable genes. **D**. Switch times can be used to rank genes by the length of priming and decoupling intervals. Each range is scaled to 1 with outliers (n=1) removed. **Top two rows**: Histogram of priming intervals. *Pbx3* and *Celsr1* possess short and long priming phases, respectively. **Bottom two rows**: Histogram of decoupled intervals. While *Rspo3* has a short decoupling phase with few cells within, *Tgfbr1*'s decoupling phase extends from RNA induction to RNA repression, and up to the end of the trajectory.

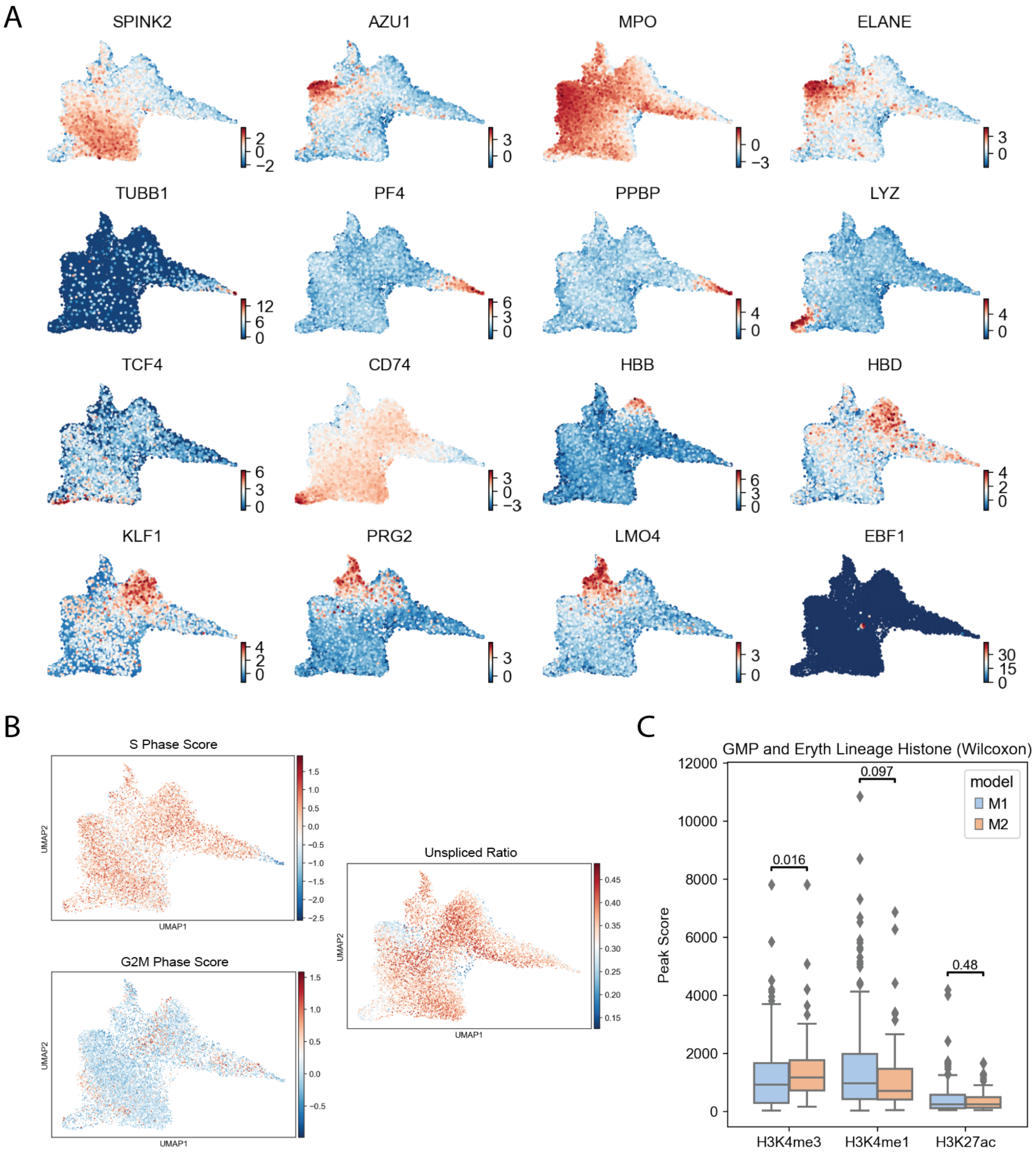


**Fig. S5. Additional figures for HSPC. A**. Canonical marker gene expression for HSPCs. **B**. Cell cycle (S phase and G2M phase) scores and total unspliced ratio (U/(U+S)) plotted on UMAP coordinates. These factors were regressed out of the total RNA expression (but not the unspliced and spliced counts) during the preprocessing step as they do not appear to be cell-type or lineage specific. **C**. Box plots of histone modification levels from bulk ChIP-seq of FACS-purified HSCs. Each point in the box plot represents the sum of histone modification signal at chromatin accessibility peaks linked to a Model 1 or Model 2 gene. P-values are from a one-sided Wilcoxon rank-sum test.

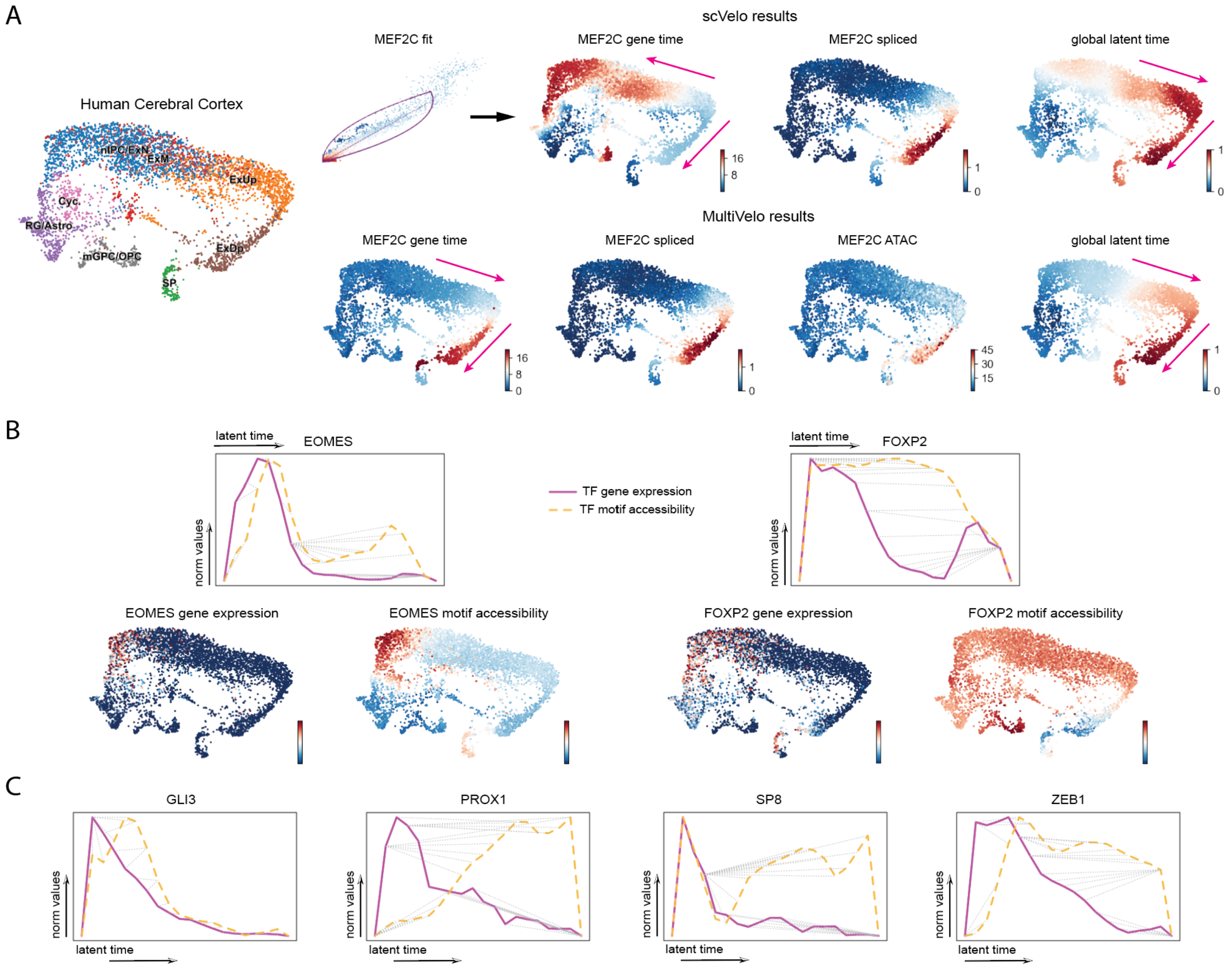


**Fig. S6. Additional figures for human brain. A**. Validation of the direction of *MEF2C*. **Left**: UMAP with cell types. **Top**: scVelo's *MEF2C* fit produces inconsistency between gene time and global latent time. **Bottom**: MultiVelo's results show consistent progression from nIPC to deeper layer (ExDp). **B**. DTW and UMAP results for *EOMES* and *FOXP2* transcription factors. **C**. Additional motif DTW alignment results showing time lags between TF gene expression and corresponding motif accessibility.
